# Human Induced Pluripotent Stem Cell based Hepatic-Modeling of Lipid metabolism associated TM6SF2 E167K variant

**DOI:** 10.1101/2023.12.18.572248

**Authors:** Lanuza AP Faccioli, Yiyue Sun, Takashi Motomura, Zhenghao Liu, Takeshi Kurihara, Zhiping Hu, Zeliha Cetin, Jonathan Franks, Donna Stolz, Alina Ostrowska, Rodrigo M Florentino, Ira J Fox, Alejandro Soto-Gutierrez

## Abstract

**BACKGROUND AND AIMS:** TM6SF2 rs58542926 (E167K) is associated with an increase in the prevalence of Metabolic Disfunction-Associated Steatotic Liver Disease (MASLD). Despite all the investigation related to the role of this variant in lipid metabolism, conflicting results in mouse studies underscore the importance of creating a human model for understanding the TM6SF2 mechanism. Therefore, the aim of this study is to generate a reliable human in vitro model that mimic the effects of the TM6SF2 E167K mutation and can be used for future mechanism studies.

**APPROACH AND RESULTS:** We performed gene editing on human-induced pluripotent stem cells (iPSC) derived from a healthy individual to obtain the cells carrying the TM6SF2 E167K mutation. After hepatic differentiation, a decrease in TM6SF2 protein expression was observed in the mutated-induced hepatocyte. An increase in intracellular lipid droplets and a decrease in the efflux of cholesterol and ApoB100 were also observed. Transcriptomics analysis showed up-regulation of genes related to the transport, flux, and oxidation of lipids, fatty acids, and cholesterol in TM6SF2 E167K cells. Additionally, signs of cellular stress were observed in the ER and mitochondria.

**CONCLUSIONS:** Our findings indicate that induced hepatocytes generated from iPSC carrying the TM6SF2 E167K recapitulate the effects observed in human hepatocytes from individuals with the TM6SF2 mutation. This study characterizes an in vitro model that can be used as a platform to help in the identification of potential clinical targets and therapies and to understand the mechanism by which the TM6SF2 E167K variant leads to vulnerability to MASLD.

## Introduction

Chronic liver disease (CLD) which produce liver failure, hepatocellular carcinoma (HCC) and portal hypertension results in 2 million deaths per year worldwide. HCC alone ranks as the 5th leading cause of cancer-related deaths in the United States (1, 2). The etiology of CLD has changed dramatically over the last few decades (3). The burden of chronic hepatitis C has diminished due to the emergence of highly effective direct-acting antiviral agents, while the incidence of Metabolic dysfunction-associated steatotic liver disease (MASLD), including Metabolic dysfunction-associated steatohepatitis (MASH), has grown. MASLD is currently the leading indication of adult liver transplantation and is the cause of CLD most associated with the development of HCC (4–8). Despite its public health importance and financial burden, there is currently no FDA-approved therapy for MASLD. The lack of therapeutic options reflects the complex patient pathogenesis and heterogeneity as well as the lack of experimental models that fully recapitulate disease phenotypes involved in the progression of MASLD. Thus, our current overall ability to predict disease progression and response to drug treatments is still limited, even among large patient cohorts.

Genome-wide association studies (GWAS) have identified numerous genetic variants (9), which are linked to medical disorders (7). Several are linked to increased vulnerability to MASLD. This include: single nucleotide polymorphisms (SNP) in PNPLA3 (rs738409 C>G p.Ie148Met; patatin-like phospholipase domain-containing protein 3), MBOAT7 (rs62641738 C>T; Membrane Bound O-Acyltransferase Domain Containing 7), GCKR (rs780094 C>T; Glucokinase Regulator) and TM6SF2 (rs58542926 C>T p.Glu167Lys; transmembrane 6 super family 2) (9–12). This last missense variant causes the substitution of glutamine with lysine at position 167 (p. Glu167Lys or E167K), and many studies indicated that this leads to misfolding of the protein, with concomitant accelerated protein degradation (10) and results in an increase in serum cholesterol levels and accumulation of lipids in the liver. Other investigators and our own studies indicate that individuals carrying the *TM6SF2* E167K variant exhibit a twofold higher prevalence of both MASH and advanced fibrosis (10, 13) compared to control subjects. In a recent study, it was reported a significant increased risk of liver transplantation/liver-related death in *TM6SF2* E167K carries (14). Because this variant is rare, its frequency in individuals of European ancestry is 7.2%, African is 3.4% and Hispano-Americans is 4.7% (15), the precise molecular mechanism by which this genetic variation produces a higher incidence of liver disease has yet to be fully elucidated (9, 10, 16).

Although there is a substantial and growing body of literature describing the role of TM6SF2 in lipid accumulation and increase in cholesterol levels, the study of this phenomenon in TM6SF2 loss-of-function or overexpression mouse models has generated inconsistent conclusions (17–19). Therefore, the creation of human model that can illuminate the effects of this variant would be of a great value for MAFLD studies and therapeutic development. Thus, we generated human induced pluripotency stem cells (iPSCs) from an individual who carried the WT *TM6SF2* allele (CC) and using CRISPR-Cas9 we generated a gene-edited iPSCs carrying the *TM6SF2* mutant allele (TT). These iPSCs lines were differentiated into hepatocytes (iHeps) (20). We found that both human iHep-TM6SF2-WT and human iHep-TM6SF2-E167K expressed hepatocyte-enriched transcription factors including the human adult isoform HNF4α, human albumin and did not express the immature marker AFP. Further characterization revealed expression differences of TM6SF2 protein, accumulation of triglycerides, Apolipoprotein B (ApoB) and total cholesterol. Additionally, we dissected the molecular consequences of *TM6SF2* E167K variant using transcriptomics and unveiled alterations related to the transport, flow, removal, and oxidation of lipids, fatty acids, and cholesterol. Collectively, our results suggest that this human model system can be used to capture and study the role of genetic variants in the development of CLD.

## Methods

### Primary human hepatocyte

Cryopreserved primary hepatocytes from healthy individuals were obtained from In Vitro ADMET Laboratories Inc. (IVAL, Columbia, MD, USA). End-stage liver disease (ESLD) hepatocytes were isolated from therapeutically resected livers and fresh human liver tissue specimens from patients (IRB: STUDY20090069) undergoing liver transplantation in the adult liver transplant programs at the University of Pittsburgh Medical Center (UPMC). Human fetal liver tissues were obtained from the University of Washington Department of Pediatrics, Division of Genetic Medicine, Laboratory of Developmental Biology (Seattle, WA) after obtaining written informed consent by a protocol approved by the Human Research Review Committee of the University of Pittsburgh (honest broker approval numbers HB015 and HB000836). Human fetal liver hepatocytes were isolated, cultured, and differentiated into fibroblasts, as previously described (21). Specific information on the age, gender, and cell viability of human liver tissue and hepatocytes used in this study is described in Supplementary Table 1 and Supplementary Table 5.

### Genotyping and Sanger sequencing

Genotyping and Sanger sequencing were performed by extracting genomic DNA with the DNeasy Blood & Tissue Kit (QIAGEN, Hilden, Germany) following the manufacturers’ instructions. Genomic DNA samples were genotyped using TaqMan SNP genotyping assays for TM6SF2 rs58542926, PNPLA3 rs738409, GCKR rs780094, MBOAT7 rs62641738, HSD17B13 rs72613567 and MTARC1 rs2642438 (ThermoFisher Scientific, Waltham, MA). Amplification and genotype clustering were performed using a StepOnePlus system (Applied Biosystems, Foster City, CA).

For sequencing, polymerase chain reaction (PCR) amplification was conducted with the KOD ONE PCR Master Mix (Toyobo, Osaka, Japan) using the forward primer (CAAGATGTCCAGCCAGAGAGG) and reverse primer (CTTTCTTGTGACAAAGGAGAACCT) for *TM6SF2*. After DNA samples were amplified, the result of the amplification was confirmed with a 2% agarose gel. PCR products were then purified using the ExoSAP-IT Express PCR cleanup kit (Applied Biosystems, Foster City, CA) and sequenced at the Genomics Research Core at the University of Pittsburgh, Pennsylvania, PA. Sequencing buffer and a 1:4 dilution of BigDye 3.1 (ThermoFisher Scientific, Waltham, MA) were added, and thermocycling was performed according to ABI recommendations. According to manufacturer instructions, removing unincorporated sequencing reagents is performed using CleanSeq magnetic beads (Agencourt, Beckman Coulter, Brea, CA). Two control samples were included with every sequencing to ensure the proper performance of reagents and equipment.

### Generation and Culture of Human iPSC

iPSC-TM6SF2-WT were generated from fibroblasts. Reprogramming of fibroblasts was performed using episomal plasmid vectors adapted from a previously described method (21). Briefly, for each nucleofection, 1 million cells were resuspended in 100 mL of the AmaxaTM NHDF Nucleofector kit (Lonza, Walkersville, MD), containing 1ug of each of the four episomal plasmid vectors encoding OCT3/4 and p53 shRNA, SOX2 and KLF4, L-MYC and LIN28, and enhanced green fluorescent protein (eGFP) (Addgene, Boston, MA, USA). Cells were nucleofected using the Amaxa 4D-Nucleofector (Lonza, Walkersville, MD) and plated in mTeSR on human embryonic stem cell-qualified Matrigel-coated plates (Corning, New York, NW).

Colonies were isolated around 60 days after induction based on morphology. The cell line underwent karyotyping, and its pluripotency was validated by the expression of NANOG, OCT4, and membrane markers SSEA and TRA-1-60 at different passages. Additionally, the cell line was routinely tested and found to be negative for mycoplasma contamination. A commercial iPS cell (WTC11) was used as a positive control (Coriell Institute, Camden, NJ).

### Gene editing

The single-guide RNA (sgRNA) sequence (GCAAATACAGCTCCGAGATC) was designed to cut the human TM6SF2 gene at position chr19:379,549 to replace the major allele (C) with the minor allele (T). The sgRNA was cloned into a plasmid vector and nucleofected into the iPSC-TM6SF2-WT together with the donor DNA (ACAGATGTCCAGCAGGGTTCTGGCATGGCTGATGCCCTCTCTCCTGCACCATGGAAGGCAAATACAGCTCCAAGATCAGACCTGCCTTCTTCCTCACCATCCCCTACCTGCTGGTGCCATGCTGGGCTGGCATGAAGGTCT), using the Amaxa 4D-Nucleofector (Lonza, Walkersville, MD). For each nucleofection, 1 million cells were resuspended in 20 uL of the Amaxa NHDF Nucleofector Kit (Lonza, Walkersville, MD), containing 1ug of each of the gRNA and donor plasmid vectors (ABM, Richmond, Canada). Antibiotic selection was performed, and DNA from selected clonal cells was extracted, amplified, and purified before sequencing as previously described. Minor homozygous clones were identified, expanded, and cryopreserved, and one clone was used to perform the experiments.

### Embryoid Body Formation

Embryoid bodies (EBs) were formed by plating iPSC-TM6SF2-WT and iPSC-TM6SF2-E167K cells at a density of 2.5×10^4^ cells per cm^2^ on low-attachment 6 well plates in mTeSR with 20% of FBS and cultured at 37°C and 5% CO_2_. The medium was changed every 72 hours. EBs started to form in suspension after one week of culture. At day 20, EBs were fixed in 4% paraformaldehyde for 24 hours and 70% ethanol overnight at 4 °C, and then embedded in paraffin. 5-micron sections were placed on glass slides and then used for immunostaining of the three germ layers.

### Quantitative Real-Time PCR

Total RNA was isolated from human cells using RNeasy Mini kits (QIAGEN, Hilden, Germany) and reverse transcribed using Super-Script III (Invitrogen, Carlsbad, CA) following the manufacturers’ instructions. We performed qPCR with a StepOnePlus system (Applied Biosystems, Foster City, CA) using TaqMan Fast Advanced Master Mix (Life Technologies, Waltham, MA). The probes used are listed in Supplementary Table 2. Relative gene expression was normalized to ß-actin (ACTB) mRNA using ΔΔCT method.

### Immunostainings

The samples were fixed with 4% PFA for 15 minutes and washed another three times with PBS. Following fixation, samples were washed 3 times with wash buffer (PBS, 0.1% BSA, and 0.1% TWEEN 20) for 5 minutes and then blocked and permeabilized in blocking buffer (PBS, 10% normal donkey or goat serum, 1% BSA, 0.1% TWEEN 20, and 0.1% Triton X-100) for 1 hour at room temperature. Subsequently, the samples were then incubated with primary antibodies in blocking buffer overnight at 4°C. The following day, samples were washed three times with wash buffer for 5 minutes and incubated with secondary antibodies in blocking buffer for 2 hours in the dark at room temperature. Then, samples were washed 3 times with wash buffer for 5 minutes, followed by 3 washes with PBS, and counterstained with 1 µg/mL of DAPI (Sigma Aldrich, ON, Canada) for 1 minute at room temperature in the dark. In the end, samples were washed three times with PBS and stored in the dark at 4°C. Samples were imaged using an Eclipse Ti inverted microscope (Nikon) and the NIS-Elements software platform (Nikon, NY, USA). Following that, images were analyzed using ImageJ software. RGB stacks were generated, pre-processed to equalize the illumination within the stack, thresholded, and measured. All antibodies used are listed in Supplementary Table 3.

To better understand the role of TM6SF2 rs58542926 in ESLD tissue and cells, we needed to validate the TM6SF2 primary antibody, as a substantial body of literature that addresses this subject shows the variability in antibodies and methodologies employed has inconsistent and frequently conflicting results (11, 17, 1). To understand the distribution of TM6SF2 in liver tissue, we analyzed ESLD tissue from patients that were WT (CC) and patients with the E167K (TT) for TM6SF2 rs58542926 (Supplementary Figure 1B). For TM6SF2 immunohistochemistry staining, 5-7-micron sections were deparaffinized with xylene and dehydrated with ethanol. Antigen unmasking was performed by boiling in 10 mM citrate buffer, pH 6.0. After the antigen unmasking, the slides were exposed to 3% hydrogen peroxide and incubated overnight at 4°C with the primary antibody. On the following day, tissue sections were incubated with the secondary biotinylated antibody corresponding to the animal species of the primary antibody (BA-1000; Vector Laboratories, Burlingame, CA) and exposed to 3,30-diaminobenzidine (SK-4105; Vector Laboratories) to visualize the peroxidase activity. Counterstaining was performed with Richard-Allan Scientific Signature Series Hematoxylin (Thermo Scientific, Waltham, MA). Samples were imaged using an Axiovert 40 CFL (Zeiss, NY, USA) microscope and the Zeiss Zen 3.8 software platform (Zeiss, NY, USA). All antibodies used are listed in Supplemental Table 3.

### Differentiation of Human iPSC into induced hepatocyte (iHep)

Our hepatocyte differentiation protocol was reported by Collin de l’Hortet,*et al*., 2019 (21). Briefly, human iPSCs were passaged with Accutase (Stem Cell Technologies, Vancouver, Canada) and re-plated at a density of 1 to 2×10^5^ per cm^2^ in growth factor reduced Matrigel (Corning Incorporated, Corning, NY) coated plates in mTeSR. The day after, cells were exposed to a defined differentiation medium containing RPMI (Invitrogen, Carlsbad, CA), 1x B-27 without insulin supplement (Invitrogen, Carlsbad, CA), 0.5% Penicillin/Streptomycin (Millipore, Billerica, MA), 0.5% non-essential amino acids (Millipore, Billerica, MA), 100 ng/mL Activin A (R&D Systems, Minneapolis, MN), 10 ng/mL BMP4 (R&D Systems, Minneapolis, MN), and 20 ng/mL FGF2 (BD, Franklin Lakes, NJ) for two days and placed in a normal O_2_ incubator (endoderm induction). Cells were subsequently maintained in a similar medium without FGF2 and BMP4 for two days in ambient O_2_ (definitive endoderm). Cells were then grown for 10 days in a defined medium containing 45% DMEM low glucose (ThermoFisher Scientific, Waltham, MA), 45% F-12 (ThermoFisher Scientific, Waltham, MA), 10% CTS Knockout SR Xenofree Medium (ThermoFisher Scientific, Waltham, MA), 0.5% Non-Essential Amino Acids (ThermoFisher Scientific, Waltham, MA), 0.5% L-glutamine (ThermoFisher Scientific, Waltham, MA), 50 ng/mL HGF (R&D Systems, Minneapolis, MN) and 1% DMSO (Sigma-Aldrich, Saint Louis, MO), The medium was changed every other day (hepatic specification).

### Enzyme-Linked Immunosorbent Assay (ELISA)

ELISA for Apolipoprotein B100 (ApoB100) was done using the ApoB100 ELISA KIT (Thermo Scientific, Waltham, MA) according to the manufacturer’s protocol. The reaction was developed for 30 minutes with 100 uL/well TMB substrate solution and stopped with 50 uL/well stop solution. HRP activity was measured in an HTX microplate reader (Biotek, Winooski, VT) at a wavelength of 450 nm. To calculate the sample value, their absorbance was interpolated with a standard curve generated using a four-parameter algorithm.

### Western Blotting

Human samples were incubated with RIPA lysis buffer (Sigma Aldrich), 1x Halt™ Protease, and Phosphatase Inhibitor Cocktail (Thermo Fisher Scientific, Waltham, MA) and incubated for 30 min at 4°C. Samples were centrifuged at 13,000g for 10 min at 4°C. The supernatant from each sample was then transferred to a new microfuge tube and used as the whole cell lysate. Protein concentrations were determined by comparison with a known concentration of bovine serum albumin using a Pierce BCA Protein Assay Kit (Thermo Fisher Scientific, Waltham, MA). 30 µg of lysate were loaded per lane into 10% Mini-PROTEAN TGX™ gel (BioRad, Hercules, CA).

Next, proteins were transferred onto the PVFD transfer membrane (Thermo Fisher Scientific, Waltham, MA). Membranes were incubated with a primary antibody solution overnight and then washed. Membranes were incubated for 1 hour in a secondary antibody solution and then washed. Target antigens were finally detected using SuperSignal™ West Pico PLUS Chemiluminescent Substrate (Thermo Fisher Scientific, Waltham, MA). Images were scanned and analyzed using ImageJ software. All band density values were normalized to the band density for GAPDH. All antibodies used are listed in Supplemental Table 3.

### Cholesterol Analysis

For analysis of cholesterol metabolism, cells were cultured for 48 hours, and the quantification of intracellular and extracellular total cholesterol and its fractions was measured using the Cholesterol Assay Kit (Abcam, Cambridge, United Kingdom) according to the manufacturer’s instructions. The fluorescence signal (Ex/Em: 535/587 nm) was measured in an HTX microplate reader (Biotek, Winooski, VT).

### Nile Red Staining

Samples were fixed with 4% PFA for 15 minutes and washed three times with PBS. After that, the samples were incubated with a 0.3 mM Nile Red (Sigma Aldrich, ON, Canada) solution for 30 minutes at room temperature. Then, they were washed twice with PBS and counterstained with 1 µg/mL of DAPI (Sigma Aldrich, ON, Canada) for 1 minute. Samples were imaged using an Eclipse Ti inverted microscope (Nikon) and the NIS-Elements software platform (Nikon, NY, USA). Following that, images were analyzed using ImageJ software.

### RNA-seq, differential gene analysis, and gene set enrichment analysis

Whole-genome strand-specific RNA-seq was used to profile RNA expression levels in iHep-TM6SF2-WT and iHep-TM6SF2-E167K. RNA-Seq libraries were prepared as previously described. RNA was extracted using TRIzol, followed by column purification (Zymo RNA Clean and Concentrator Column) according to the manufacturers’ instructions. Total RNA was depleted of ribosomal RNA using pooled antisense oligo hybridization and depletion through RNaseH digestion. Following purification over a Zymo RNA clean and concentrator column, first-strand complementary DNA (cDNA) was synthesized. Subsequently, second-strand cDNA was synthesized, purified, and fragmented. RNA-seq libraries were prepared using Illumina technology (Illumina DRAGEN RNA, 3.10.12). Briefly, end repair, A-tailing, and barcoded adapter ligation were followed by polymerase chain reaction amplification and size selection. The integrity of the libraries was confirmed by quBit quantification, fragment analyzer size distribution assessment, and Sanger sequencing of about 10 fragments from each library. Libraries were sequenced using paired-end Illumina sequencing.

RNA-seq data were proceeded for DEG using the R software (v.4.2.3) DESeq2 package. The adjusted *p*-value cutoff was set at 0.05, and logFC > 1.5 was the filter criteria. The GSEA analysis was done using the R package WebGestaltR package and GSEA software (Gene Set Enrichment Analysis, v4.3.2) from the Broad Institute. Gene collections were obtained from the MSigDB KEGG subset of CP. These results were further visualized with R software (v.4.2.3) and package ggplot2 (v.3.4.2).

### Transcription Profiling by the RT2 Profiler PCR Array

Total RNA was isolated using the RNeasy Mini kit (QIAGEN, Hilden, Germany) and reverse transcribed using SuperScript III (Invitrogen, Carlsbad, CA) following the manufacturers’ instructions, and complementary DNA was amplified. Key genes involved in the regulation and enzymatic pathways of fatty liver were simultaneously assayed with the RT2 Profiler PCR Array Human Fatty Liver (PAHS-157ZC-6) (QIAGEN, Hilden, Germany) according to the manufacturer’s instructions and analyzed with the Data Analysis Center (QIAGEN Hilden, Germany). Ingenuity pathway analysis (IPA) was used to identify differentially expressed genes, predict downstream effects, and identify targets (QIAGEN Bioinformatics; www.qiagen.com/ingenuity). Regulatory effects analysis within IPA was used to identify the relationships between upstream regulators and biological functions.

### Transmission electron microscopy

Human samples were briefly centrifuged and washed with a PBS solution. Samples were then fixed with 2.5% glutaraldehyde overnight at 4°C. Fixed samples were processed by the Center for Biologic Imaging at the University of Pittsburgh and treated with 1% osmium tetroxide and 1% potassium ferricyanide for 1 hour at room temperature. Samples were washed with PBS and dehydrated in a graded series of ethanol solutions (30%, 50%, 70%, and 90%—10 minutes each) and three 15-minute changes in fresh 100% ethanol. Infiltration was done with four 1-hour changes of EPON embedding plastic. The last change of EPON was allowed to polymerize overnight at 37°C and then for 48 hours at 60°C. Resin blocks were removed from the Eppendorf tubes, and 70 nm sections were taken and placed onto copper TEM grids. Image acquisition was made using either the JEM-1011 or the JEM-1400Plus transmission electron microscopes (Jeol, Peabody, MA) at 80kV fitted with a side mount AMT 2k digital camera (Advanced Microscopy Techniques, Danvers, MA).

### Statistical Analysis

For statistical analysis, means between two groups were compared by the *t* test. Since data for continuous variables were not normally distributed, p-values (p) were determined using an unpaired, two-tailed Welch’s t-test with 95% confidence. Data are reported as means ± SD, and p-values ≤0.05 were considered statistically significant. Statistical analyses were performed using GraphPad Prism version 9.3.0.

## Results

### Generation of a human iPSC and edition of the TM6SF2-E167K variant

First, we assessed the impact of TM6SF2 rs58542926 C>T p.Glu167Lys frequency in healthy subjects and ESLD patients in a United States cohort (13). We focused on TM6SF2 rs58542926 C>T variant in donors and explanted human cirrhotic livers with end-stage liver disease (ESLD) due to metabolic dysfunction-associated steatohepatitis (MASH) where robust GWAS have linked this variant to a spectrum of liver diseases ranging from steatosis to metabolic dysfunction-associated steatohepatitis (MASH), hepatic fibrosis, ESLD, HCC and increased risk of mortality in the general population. The TM6SF2-E167K gene variant was present in less than 1% of healthy individuals, and 1.8% of patients with MASH-associated ESLD (Figure 1A and Supplementary table 5). This analysis show that TM6SF2-E167K gene variant was present at a low frequency in this small (healthy donors n=123; ESLD n=50) cohort analyzed (13) and its frequency was maintain in ESLD (16,17), thus, demonstrating the need to generate alternative human liver models to understand the role of TM6SF2-E167K gene variant in the development of CLD.

**Figure 1:**
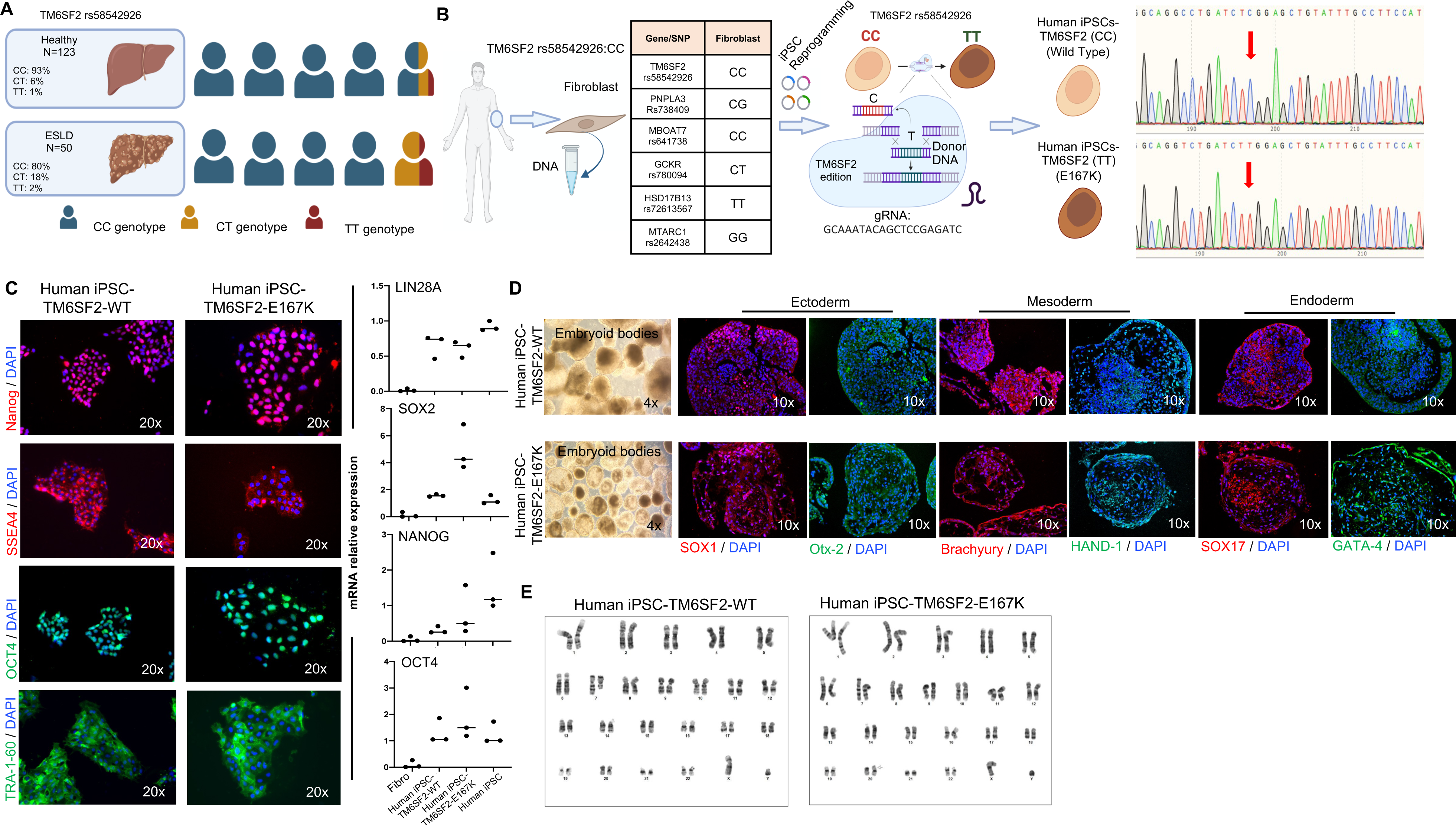
Generation and characterization of iPSC-TM6SF2-WT and iPSC-TM6SF2-E167K. (A) Genotype frequency of the *TM6SF2* rs58542926 variant in a US cohort (healthy individuals, n = 123, and ESLD samples, n = 50). Human symbols represent 20% of the prevalence. (B) Schematic design of the generation of iPSC-TM6SF2-WT and iPSC-TM6SF2-E167K. We first generated iPSC-TM6SF2-WT from fibroblasts obtained from a healthy individual. Followed by gene editing using CRISPR/Cas9 to generate iPSC-TM6SF2-E167K. The Sanger sequence for the *TM6SF2* gene confirmed that iPSC-TM6SF2-WT cells are major homozygous (CC) and iPSC-TM6SF2-E167K cells are minor homozygous (TT) after gene editing for the TM6SF2 rs58542926, as indicated by the red arrow. (C) Immunofluorescence micrographs of pluripotency markers: Nanog, SSEA4, OCT4, and TRA-1-60 (left panel) and quantitative gene expressions of pluripotency markers: SOX2, LIN28A, OCT4, and Nanog (right panel) in both iPSC-TM6SF2-WT (n = 3) and iPSC-TM6SF2-E167K (n = 3). WTC11 cells were used as a positive control, and human fibroblasts were used as a negative control. Values are determined relative to β-actin and presented as fold change relative to the expression in human WTC11, which is set as 1. (D) Micrographs of embryoid bodies and immunofluorescence micrographs of the three germ layer markers: ectoderm (SOX1, OTX-2), mesoderm (HAND-1, Brachyury), and endoderm (SOX17, GATA-4) in both iPSC-TM6SF2-WT and iPSC-TM6SF2-E167K. (E) G-banding analysis for karyotype in both iPSC-TM6SF2-WT and iPSC-TM6SF2-E167K shows no abnormalities in the cells.

Then, we identified human fibroblast carrying the major allele TM6SF2 rs58542926:C while other variants predictive of liver disease (MBOAT7 rs641738, TM6SF2 rs58542926, and GCKR rs780094, HSD17B13 rs72613567, MTARC1 rs2642438) were not present (Figure 1B). After genotyping screening, human fibroblast was reprogrammed into iPSCs as previously described (20, 21). The resulting human iPSC line (iPSC-TM6SF2-WT) was then single nucleotide edited to carry TM6SF2-E167K gene variant CRISPR-Cas9 mediated edition, we named these cells iPSC-TM6SF2-E167K. The resulting human iPSCs (iPSC-TM6SF2-WT and iPSC-TM6SF2-E167K) were cultured for >10 passages before a complete characterization and validation studies were performed. Successful single base editing was confirmed by sanger sequencing and showed the presence of the gene variant for *TM6SF2* rs58542926 C>T (Figure 1B).

Human iPSC-TM6SF2-WT and iPSC-TM6SF2-E167K showed normal pluripotent morphology, consisting of compact colonies with distinct borders like that seen in human embryonic stem cells (hESCs), expressed NANOG, SSEA4, OCT4 and TRA-1-60, and exhibited similar mRNA expression of pluripotency markers (Lin28A, SOX2, Nanog, and OCT4) comparable to that seen in control human iPSCs (Figure 1C). Embryoid Bodies (EB) derived from Human iPSC-TM6SF2-WT and iPSC-TM6SF2-E167K lines formed all three germ layers (Figure 1D) which was corroborated by the spontaneous expression of ectodermal (SOX1 and Otx-2), mesodermal (Brachyury and HAND-1), and endodermal (SOX17 and GATA-4) markers (Figure 1D). Both human iPSC-TM6SF2-WT and iPSC-TM6SF2-E167K cells also had a normal karyotype (Figure 1E).

### Hepatocyte-Directed Differentiation of Human iPSC-TM6SF2-WT and iPSC-TM6SF2-E167K

We then proceed to differentiate the human iPSC-TM6SF2-WT and iPSC-TM6SF2-E167K toward hepatocytes using our previously published protocol (20–23). Cells were first cultured with a combination of activin A, bone morphogenetic protein 4 (BMP4), and fibroblast growth factor (FGF)-2 to induce definitive endoderm. After showing the presence of the endoderm marker (SOX17) (Figure 2B), cells were cultured for 10 days in the presence of dimethyl sulfoxide (DMSO) and human hepatocyte growth factor (hHGF) to induce hepatocyte specificity (Figure 2A). Following the hepatocyte-directed differentiation, both human iPSC-TM6SF2-WT and iPSC-TM6SF2-E167K lines developed characteristics of hepatocytes as assessed by immunofluorescence, for the adult isoform of hepatocyte nuclear factor 4 alfa (HNF4α) and human albumin (ALB). Expression of AFP (alpha-fetoprotein), an immature hepatocyte marker, was not observed (Figure 2B). In addition, both hepatocyte differentiated cell lines (human iHeps-TM6SF2-WT and iHeps-TM6SF2-E167K) expressed critical hepatocyte-specific transcripts: HNF4α, forkhead box protein A2 (FOXA2), forkhead box protein A1 (FOXA1), hepatocyte nuclear factor 1 alpha (HNF1α), CCAAT Enhancer Binding Protein Alpha (CEBPA), retinoid X receptor (RXR), liver X receptor (LXR), peroxisome proliferator-activated receptor alpha (PPARa), sterol regulatory element-binding transcription factor 1 (srebp1c), acetyl-CoA carboxylase (ACC), fatty acid synthase (FASN) and epidermal growth factor receptor (EGFR). All were expressed at levels comparable to those found in human adult hepatocytes (Figure 2C).

**Figure 2:**
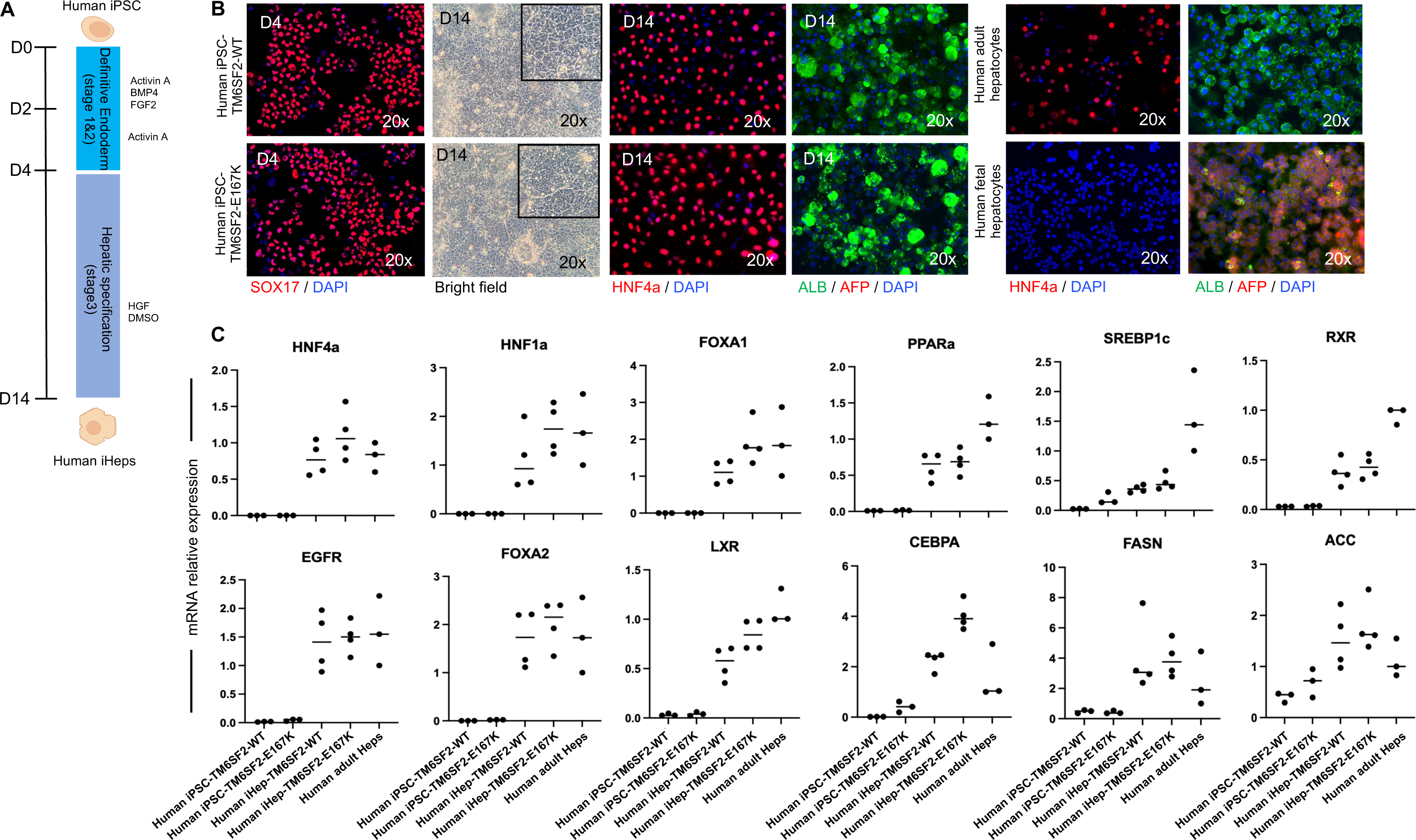
Hepatic differentiation and characterization of iHep-TM6SF2-WT and iHep-TM6SF2-E167K. (A) schematic illustration of the hepatocyte differentiation protocol, highlighting the three main stages of differentiation by sequential addition of defined medium protocols containing Activin-A, BMP-4, and FGF2 (stage 1); Activin-A (stage 2); and dimethyl sulfoxide (DMSO) and hepatocyte growth factor (HGF) (stage 3). (B) Immunofluorescence micrographs (left panel) of endoderm marker SOX17 in both iDE-TM6SF2-WT (n = 4) and iDE-TM6SF2-E167K (n = 4). Bright field micrographs of each iHep-TM6SF2-WT and iHep-TM6SF2-E167K show the cells on the last day of stage 3 and immunofluorescence micrographs of hepatocyte markers, adult isoform HNF4a, AFP, and albumin in both iHep-TM6SF2-WT and iHep-TM6SF2-E167K. Human adult hepatocyte (PHH) (n = 3) and human fetal hepatocyte (n = 3) were used as positive and negative controls, respectively. (C) Quantitative gene expression for hepatocyte markers: HNF4a, HNF1a, FOXA1, FOXA2, PPARα, LXR, RXR, FASN, EGFR, SREBP1c, ACC, and CEBPA in both iHep-TM6SF2-WT (n = 4) and iHep-TM6SF2-E167K (n = 4). PHH cells (n = 3) were used as a positive control, and both undifferentiated iPSC cells (n = 3) were used as a negative control. Values are determined relative to β-actin and presented as fold change relative to the expression in PHH, which is set as 1.

### TM6SF2-E167K variant induces protein loss-of-function and modifies lipid metabolism in human iHeps

We next studied TM6SF2 transcript and protein expression by qPCR, immunofluorescence, and western blot. We found that TM6SF2 transcript levels were not significantly different within human iHeps-TM6SF2-WT, iHeps-TM6SF2-E167K, human liver tissue or isolated human adult ESLD hepatocytes (Figure 3A). However, when protein expression was analyzed, iHeps-TM6SF2-E167K showed a significant reduced expression when compared to control human iHeps-TM6SF2-WT (Figure 3A). A hallmark of MASLD progression especially in individuals carrying the TM6SF2-E167K variant is disruption of lipid and cholesterol regulation (10, 24). Thus, we next investigate the effect of TM6SF2-E167K variant on intracellular and extracellular lipid accumulation in iHeps by performing Nile red staining and observed a significant increase in the concentration of intracellular lipids especially in iHep-TM6SF2-E167K when compared with iHep-TM6SF2-WT controls (Figure 3B). This prompt us to examine first the expression of elongation of very-long-chain fatty acid protein 6 (ELOVL6), a critical enzyme involved in lipid synthesis. We found that ELOVL6 transcript expression was significantly increased in iHeps-TM6SF2-E167K when compared to control iHeps-TM6SF2-WT (Figure 3B). Next, we studied whether the TM6SF2-E167K variant would affect cholesterol transporters such as ApoB100 and found that iHeps-TM6SF2-E167K intracellular ApoB100 protein expression was significantly increased while extracellular secretion of ApoB100 was significantly reduced when compared to iHep-TM6SF2-WT controls (Figure 3C). To confirm these findings, we measured intracellular total cholesterol and found significant increase of intracellular total cholesterol in iHeps-TM6SF2-E167K when compared to when compared to iHep-TM6SF2-WT controls. However, when we analyzed other lipoprotein transporters such HDL, while a noticeable tendency was evident, no difference was observed in the content of HDL between human iHeps-TM6SF2-WT and iHeps-TM6SF2-E167K (Figure 3D).

**Figure 3:**
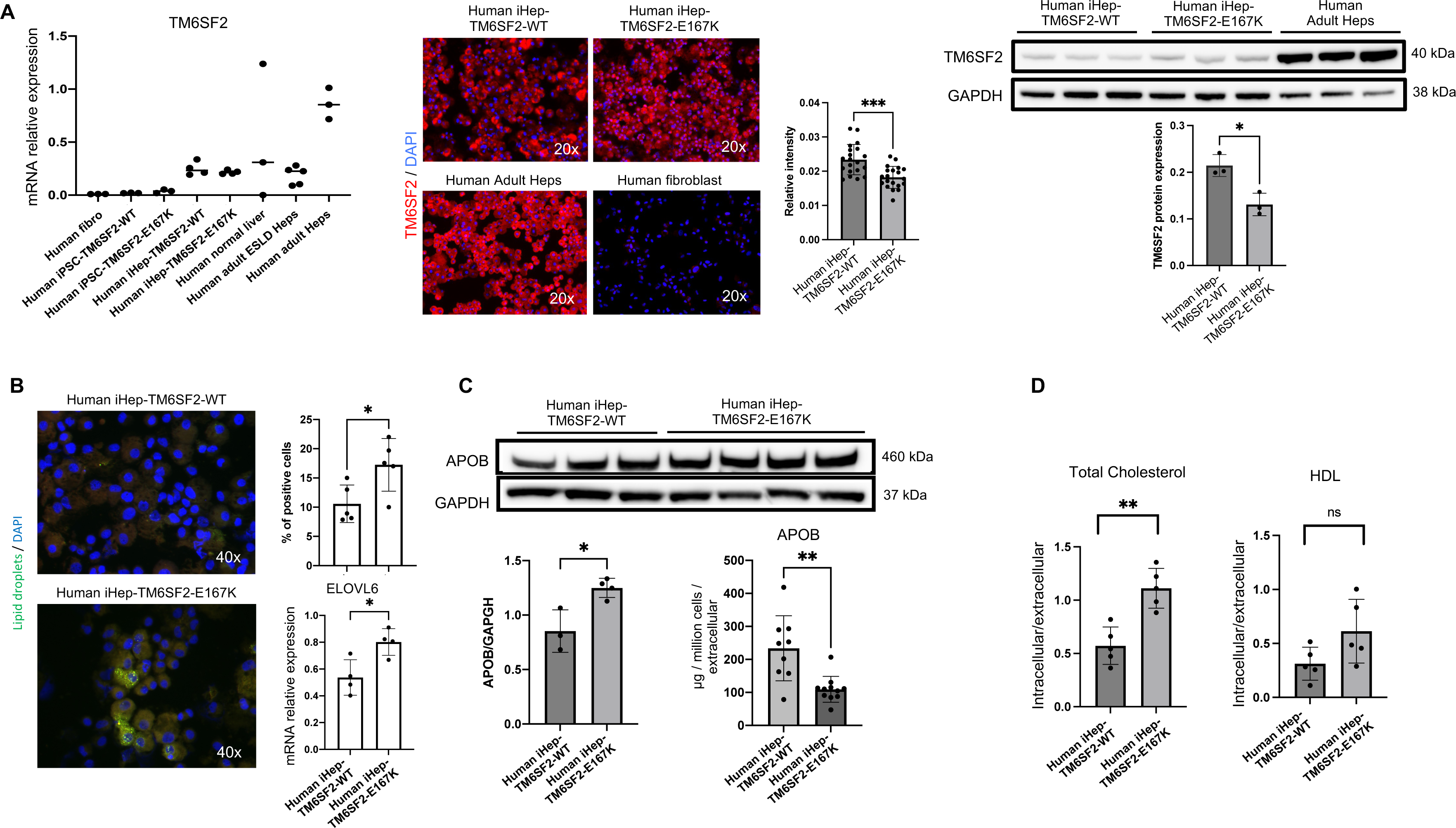
The TM6SF2 E167K mutation results in loss of function and alters lipid metabolism in human iHep. (A)*TM6SF2* expression in both iHep-TM6SF2-WT and iHep-TM6SF2-E167K. The left upper panel shows quantitative gene expression. iPSC-TM6SF2-WT (n = 3), iPSC-TM6SF2-E167K (n = 3), and human fibroblast cells (n = 3) were applied as negative controls. Human normal liver tissue (n = 3), human ESLD-WT hepatocytes (n = 5), and adult PHH (n = 3) were used as positive controls. Values are determined relative to β-actin and presented as fold change relative to the expression in PHH, which is set as 1. The middle upper panel shows immunofluorescence micrographs of the TM6SF2 marker in iHep-TM6SF2-WT and iHep-TM6SF2-E167K. Adult PHH was used as a positive control and fibroblast as a negative control. The relative TM6SF2 intensity showed a significant decrease in iHep-TM6SF2-E167K cells when compared to iHep-TM6SF2-WT (mean ± SD ***p = 0.0002 unpaired Welch’s t-test, n = 20 cells). The same was observed by the western blot. The bar chart shows the quantification of the protein expression. There was a significant decrease in iHep-TM6SF2-E167K in comparison to iHep-TM6SF2-WT (mean ± SD *p = 0.0129, unpaired Welch’s t-test, n = 3). (B) Nile red staining micrographs in both iHep-TM6SF2-WT and iHep-TM6SF2-E167K show that iHep-TM6SF2-E167K has a higher intracellular lipid droplet content when compared to iHep-TM6SF2-WT. Quantification shows a significant increase in the percentage of Nile red signal when the cells carry the E167K mutation (mean ± SD *p = 0.0274, unpaired Welch’s t-test n = 5). iHep-TM6SF2-E167K showed an increase in expression of *ELOVL6* when compared to iHep-TM6SF2-WT (mean ± SD *p = 0.0430, unpaired Welch’s t-test, n = 4). Values are determined relative to β-actin. (C) ApoB100 secretion is impaired in iHep-TM6SF2-E167K. The intracellular content of ApoB100 in iHep-TM6SF2-WT and iHep-TM6SF2-E167K was quantified by western blot. The bar charts show an increase of ApoB100 inside the iHep-TM6SF2-E167K (mean ± SD *p=0.0144, unpaired Welch’s t-test), iHep-TM6SF2-WT (n = 3), and iHep-TM6SF2-E167K (n = 4). The secretion of ApoB100 in iHep-TM6SF2-WT and iHep-TM6SF2-E167K was evaluated by ELISA and showed a decrease of this apolipoprotein in iHep-TM6SF2-E167K (mean ± SD **p=0.0042, unpaired Welch’s t-test) in iHep-TM6SF2-WT (n = 9) and iHep-TM6SF2-E167K (n = 11). (D) Intracellular total cholesterol and HLD amounts were measured in iHep-TM6SF2-WT and iHep-TM6SF2-E167K. The bar charts show a significant increase in the intracellular and extracellular ratio of total cholesterol in iHep-TM6SF2-E167K when compared to iHep-TM6SF2-WT (mean ± SD **p=0.0079, unpaired Welch’s t-test, iHep-TM6SF2-WT, n = 4 and iHep-TM6SF2-E167K, n = 4).

### Global transcriptomic characterization of iHeps-TM6SF2-E167K revealed modified lipid metabolism and cellular stress

Human livers undergo profound transcriptional and metabolic changes throughout the development of disease such as MASLD, MASH and cirrhosis and TM6SF2-E167K appears to influence disease development (13). To further characterize these changes, we analyzed the transcriptomic signature using RNA-seq. After the differential expression analysis, the transcripts indicated an up-regulation of 153 genes and down-regulation of 267 genes (Figure 4A and Supplementary Table 4). Pathway enrichment analysis indicated that there was an increase in the expression of genes related to cholesterol, fatty acid (FA), and glucose metabolism in our iHep-TM6SF2-E167K cells, when compared to iHep-TM6SF2-WT cells. The pathways up-regulated observed in gene set enrichment analysis (GSEA) are listed in Figure 4A.

**Figure 4:**
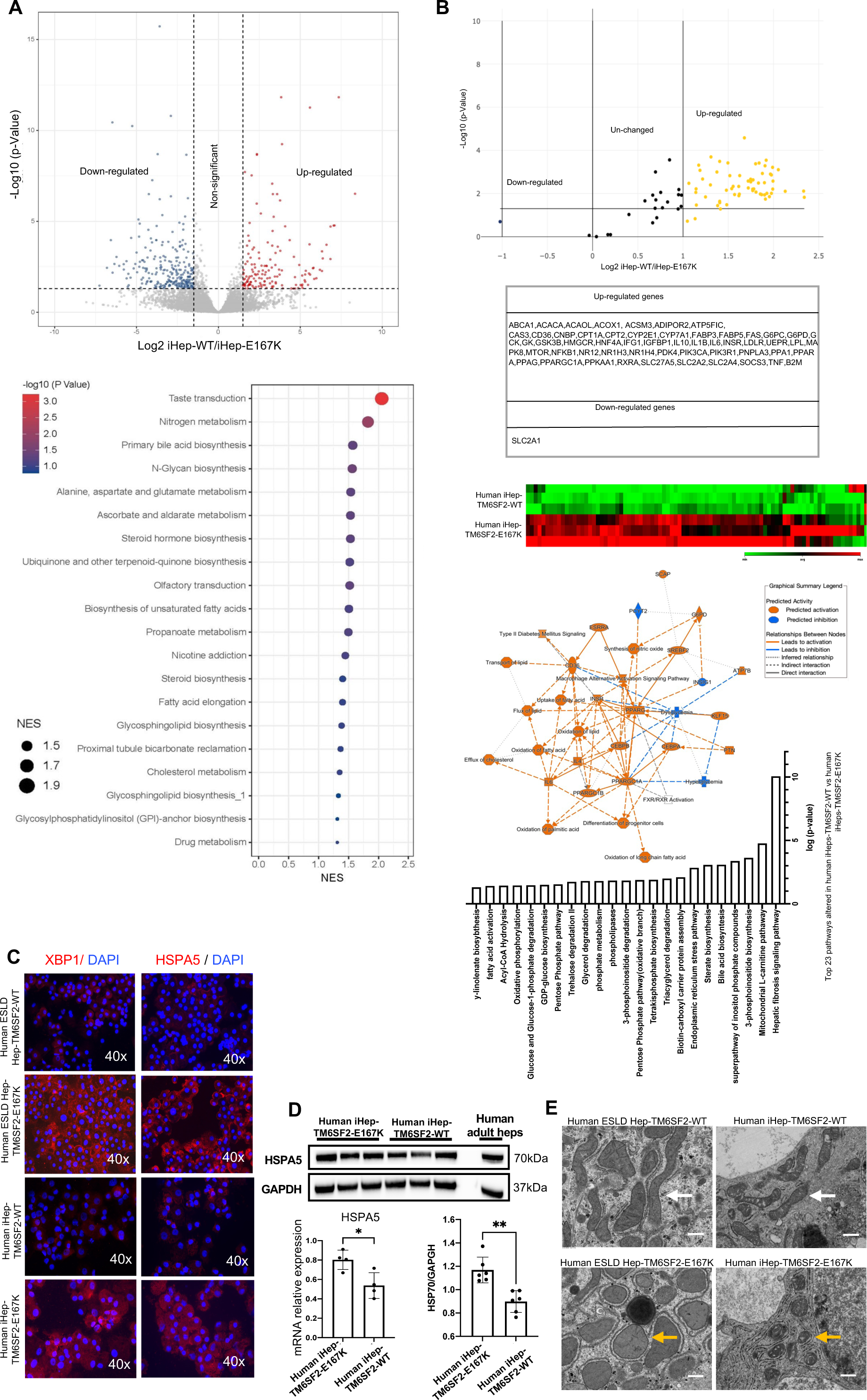
Transcription profiling analysis of iHep-TM6SF2-WT and iHep-TM6SF2-E167K and signs of cellular stress in the TM6SF2 E167K mutated cells. (A) Volcano plot showing the differential gene expression analysis of three independent differentiations of iHep-TM6SF2-WT and four independent differentiations of iHep-TM6SF2-E167K (left upper); the blue dots represent the down-regulated genes, and the red dots represent the up-regulated genes. The cut-off value for log_2_FC is 1.5 (adjust *p =* 0.05). The GSEA showed up-regulated signaling pathways related to the TM6SF2-E167K mutation. (B) Volcano plot (upper panel) and heatmap (lower panel) showing human fatty liver metabolism RNA array analysis of three independent differentiations of iHep-TM6SF2-WT and iHep-TM6SF2-E167K. The table shows the up- and down-regulated genes. Pathway analysis is based on the results from fatty liver metabolism RNA array analysis and the relationships between upstream regulators and biological functions. The top 23 pathways related to the iHep-TM6SF2-E167K mutation were ranked based on their p-values. (C) Immunofluorescence micrographs of XBP1 and HSPA5, proteins related to ER stress, in ESLD-TM6SF2-WT (n = 1), ESLD-TM6SF2-E167K (n = 1), iHep-TM6SF2-WT (n = 3), and iHep-TM6SF2-E167K (n = 3), showing an increase of these proteins in the mutant samples (ESLD-TM6SF2-E167K and iHep-TM6SF2-E167K). (D) The relative HSPA5 expression showed a significant decrease in iHep-TM6SF2-E167K when compared to iHep-TM6SF2-WT (mean ± SD*p=0.0188, unpaired Welch’s t-test n = 3). Values are determined relative to β-actin. The same was observed by the western blot. The bar chart shows the quantification of HSPA5 protein expression, and a significant decrease was observed in iHep-TM6SF2-E167K in comparison to iHep-TM6SF2-WT (mean ± SD **p=0.0010, unpaired Welch’s t-test n = 3). (E) Transmission electron microscopy (TEM) images of ESLD-TM6SF2-WT (n = 1), ESLD-TM6SF2-E167K (n = 1), iHep-TM6SF2-WT (n = 3), and iHep-TM6SF2-E167K (n = 3). The white arrows indicate the rod mitochondrial shape commonly found in human hepatocytes. The yellow array shows the round mitochondrial shape in hepatocytes that carried the TM6SF2 E167K mutation (scale bar: 600nm).

To further corroborate the transcriptomics analysis, we performed a focused RNA array. Between 56 genes that were up-regulated, we observed an increase in the expression of genes associated with fatty acid oxidation (PPARGC1A, ACOX1, ACSM3), uptake of fatty acids (CD36), lipid transportation (CPT1A, CPT2) and glucose metabolism (SLC2A2, SLC2A4) in iHep-TM6SF2-E167K, when compared to iHep-TM6SF2-WT (Figure 4B). Further pathway analyses unveiled elevated activities related to the transport, flow, removal, and oxidation of lipids, fatty acids, and cholesterol (Figure 4B). Further pathway analyses unveiled elevated activities related to the transport, flow, removal, and oxidation of lipids, fatty acids, and cholesterol (Figure 4B). Furthermore, it has been reported that excessive accumulation of lipids in hepatocytes, can lead to ER stress (25). This occurs when the ER is unable to properly process and fold proteins or metabolize lipids, resulting in a cellular stress (26). Thus, we analyzed genes that hold significance for ER and mitochondrial stress such as Heat Shock Protein Family A (Hsp70) Member 5 (HSPA5), a chaperone protein in the ER that helps fold and assemble proteins which is highly express in ER stress (27) and X-box binding protein 1 (XBP1), a transcription factor that plays a key role in ER stress (28). Both HSPA5 and XBP1 were evaluated by immunofluorescence (Figure 4C). We found that both proteins were increase in iHep-TM6SF2-E167K at levels observed in human hepatocytes freshly isolated from livers with ESLD and carrying the TM6SF2-E167K variant (Figure 4C). This observation was also corroborated at mRNA and protein levels using qPCR and western blot for HSPA5 (Figure 4D). Finally, we found an increased number of spherical mitochondria (yellow arrow) among Human hepatocytes ESLD-E167K and iHep-TM6SF2-E167K when compared to controls Human hepatocytes ESLD-WT and iHep-TM6SF2-WT (Figure 4E). Taken together, these data demonstrate that alterations in the lipid synthesis driven by TM6SF2-E167K variant can increase the susceptibility ER stress.

## Discussion

There is increasing evidence that *TM6SF2* rs58542926 plays a significant role in the metabolic processing of hepatic lipids, particularly when these lipids accumulate to excessive levels. Lipotoxicity within the liver can trigger inflammation, oxidative stress, and cellular injury, ultimately contributing to the development of MASLD (29, 30). MAFLD is marked by an abundance of fat accumulating in the liver and hereditary component (31). MASLD encompasses a spectrum of conditions, ranging from simple fat accumulation (steatosis) to more severe disorders like MASH (32).

Since this allele is relative rare and is difficult to study primary tissue, researchers have invested in animal studies. However, that mouse models don’t model the effects of this gene like in humans. Moreover, the mouse and human (33) TM6SF2 proteins are only approximately 78% identical. Therefore, we generated iHep-TM6SF2-E167K in the hope of create a model to study the impact of *TM6SF2* rs58542926 in MASH. In this study, we generated iPSC from a healthy individual (WT), followed by gene edition (22). Induced hepatocytes obtained from these cells were with upregulation of total cholesterol, intracellular content of ApoB100 (and lower secretion), intracellular fat content, ER and mitochondria stress markers and fatty acid biosynthesis pathways.

Is already described the negative effect of lipid accumulation inside the cells (34, 35). We observed an increase in the lipid droplets inside iHep-TM6SF2-E167K when compared to iHep-TM6SF2-WT. This result corroborates with our finds in mRNA levels of ELOVL6 enzyme. The enzyme ELOVL6, situated in the endoplasmic reticulum, plays a pivotal role in extending the chains of saturated and monounsaturated fatty acids (FAs). These elongated long-chain FAs constitute essential components of phospholipids, sphingolipids, cholesterol esters, and triglycerides (31). Prior investigations reveal a positive correlation between ELOVL6 expression and the severity of hepatosteatosis and liver damage in patients with MASH (36). Furthermore, it serves as a critical modulator, influencing inflammatory responses, oxidative stress, and fibrosis induced by an atherogenic high-fat diet in the liver. We observe a significant increase of this gene expression in mRNA levels in iHep-TM6SF2-E167K, indicating that these cells were with higher cellular stress.

We also noted a heightened presence of genes and protein levels associated with mitochondrial stress in iHep-TM6SF2-E167K, indicating potential overall cellular damage (37, 38). It is well established that HSP70, a crucial protein associated with mitochondrial and ER stress, plays a vital role in the generation, proper folding, and transportation of misfolded proteins to proteolytic enzymes within the mitochondrial matrix. Moreover, we detected irregularities in mitochondrial structure, as evidenced by transmission electron microscopy, aligning with the characteristics of ER stress, as previously described by Dixon *et al*. in 2012 (39). Earlier studies have established a connection between mitochondrial stress and metabolic disorders such as hepatic steatosis, insulin resistance, and type 2 diabetes (40). All these results show overall cellular damage, corroborates with the upregulation of genes involved in cholesterol, fatty acid (FA), and glucose metabolism in our iHep-TM6SF2-E167K cells. Pathway analyses revealed heightened activities in the transport, flux, efflux, and oxidation of lipids, FA, and cholesterol.

Finally, we analyzed both intracellular and extracellular ApoB100 and cholesterol levels. ApoB100 is a protein primarily associated with lipoproteins. It plays a critical role in lipid metabolism and transport in the body, specifically in the transport of cholesterol and triglycerides. Dysregulation of ApoB100 production and metabolism can lead to lipid disorders such as hyperlipidemia and contribute to conditions like MALFD. Our cholesterol results revealed a significant difference in total cholesterol, with a notable variation in the ratio between intracellular and extracellular content. In terms of ApoB100, our findings demonstrated a significant decrease in ApoB100 secretion in iHep-TM6SF2-E167K, while conversely, there was a higher ApoB100 intracellular content. In 2019, Prill *et al*. demonstrated that having only one allele copy (CT) of the *TM6SF2* E167K mutant protein is linked to the upregulation of cholesterol and fatty acid biosynthesis pathways, along with decreased ApoB100 secretion in 3D spheroid cultures of primary human mutant hepatocytes (24). Some diverge results were found in animal models also, indicating that the quantity of ApoB100 particles secreted by *TM6SF2* KO mice remained unchanged (18). These conflicting findings raise questions about the extent to which results from mouse models can be extrapolated to human physiology, emphasizing the evident need for clinical metabolic research to address this issue (41, 42).

Therefore, the iHep-TM6SF2-E167K will be useful for further investigation of the role of *TM6SF2* rs58542926 in MASH patients and will help in the identification of the mechanism by which this variant increases the risk of liver disease. The biggest limitation of our study is the small number of human hepatocytes derived from individuals carrying the *TM6SF2* E167K to use as control, due to the relatively low frequency of the E167K TT allele.

## Author Contributions

Methodology, Investigation, Validation, Data Curation, Writing - Original Draft, Review & Editing LAPF; Methodology, Investigation, Validation, Data Curation, YS; Methodology, Investigation, Validation, Data curation TM, acquisition of data, ZL; Acquisition of data, TK; Acquisition of data, ZH; Acquisition of data, ZC; Acquisition of data, JF, Acquisition of data, DS; Acquisition of data, AO, Supervision, Methodology, RMF; Conceptualization, Writing - Review & Editing, IJF; Conceptualization, Methodology, Funding acquisition, Resources, Project administration, Writing - Review & Editing, AS-G.

## Competing interest statement

A.S.-G., and I.J.F., are co-founders and have a financial interest in Von Baer Wolff, Inc. a company focused on biofabrication of autologous human hepatocytes from stem cells technology. A.O., I.J.F. and A.S.-G., are co-founders and have a financial interest in Pittsburgh ReLiver Inc, a company focused on reprogramming liver failure and their interests are managed by the Conflict-of-Interest Office at the University of Pittsburgh in accordance with their policies.

## Financial Disclosure

This work was supported by NIH grants DK099257, TR003289, DK096990, DK117881, DK119973, TR002383 to A.S.-G. This work was also supported by NIH grant 1P30DK120531- 01 to the Human Synthetic Liver Biology Core and the Pittsburgh Liver Research Center (PLRC) and internal funds from the Center for Transcriptional Medicine at the University of Pittsburgh.

---

**Supplementary Figure 1:**
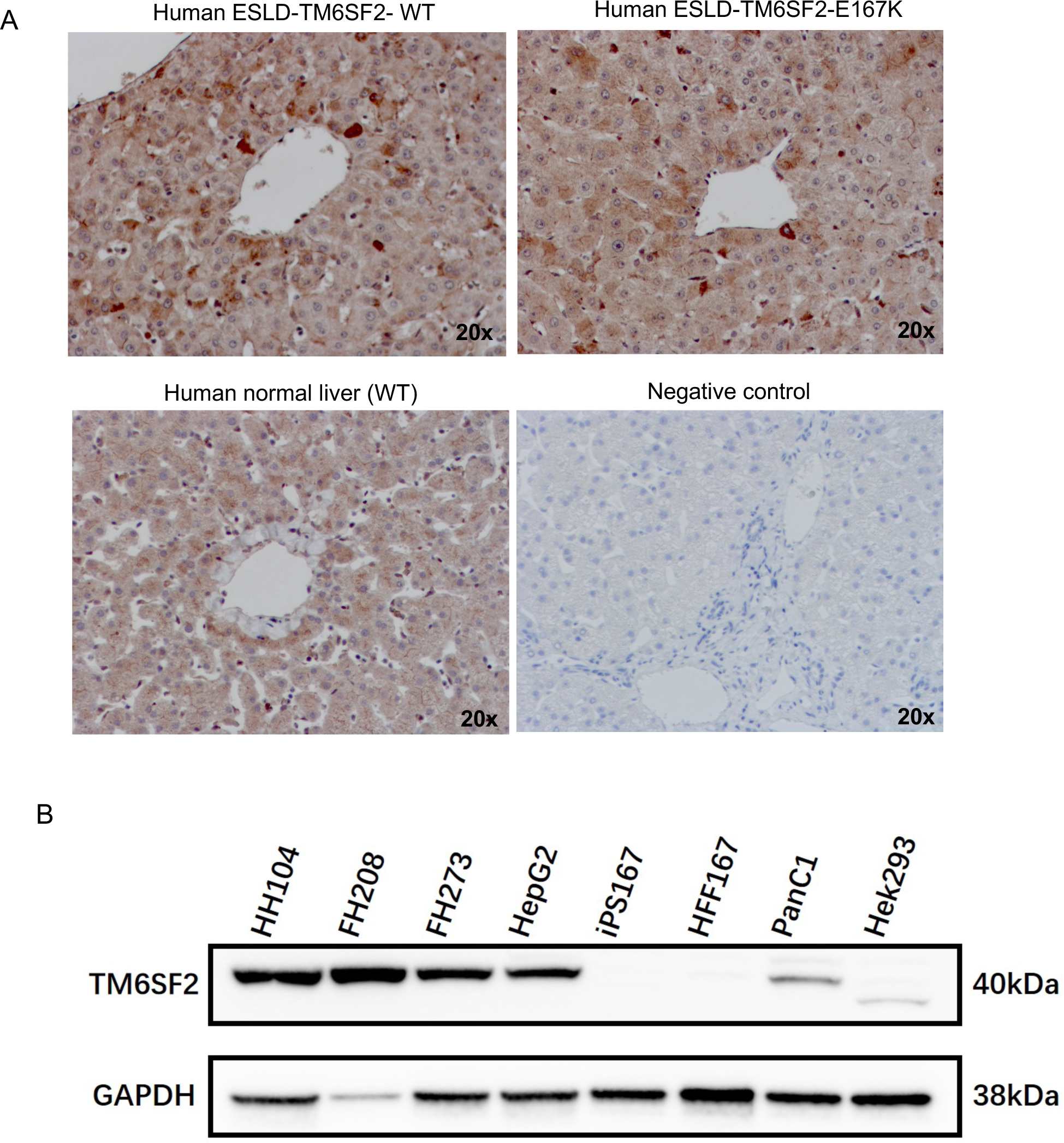
TM6SF2 antibody validation. (A) Micrographs of immunohistochemistry staining of TM6SF2 in human ESLD TM6SF2-WT tissue (n = 1) and human ESLD TM6SF2-E167K tissue (n = 1) compared to human normal liver tissue carrying TM6SF2-WT (n = 3). (B) Western blot showing that TM6SF2 is present in primary human hepatocytes (HH104), fetal human hepatocytes (FH208 and FH 273), HepG2 cells, Panc1, and Hek293 and not observed in iPSC and human fetal fibroblasts (HFF167). GAPDH was used as a loading control.

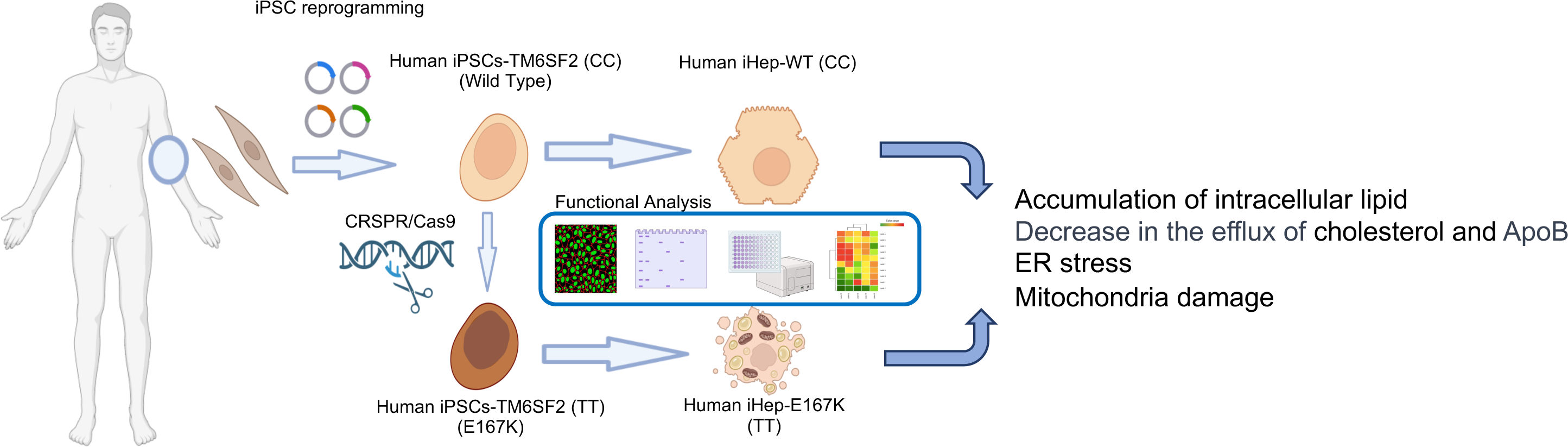

