## Supplementary Table 4 for "Human Induced Pluripotent Stem Cell based Hepatic-Modeling of Lipid metabolism associated TM6SF2 E167K variant"

Supplementary table 4: Up and down regulated genes analysed by RNA-seq

| Up-regulted genes | | | | | | |
|---|---|---|---|---|---|---|
| Gene | BaseMean | log2FoldChange | lfcSE | stat | pvalue | padj |
| ENSG00000010379.16 | 134.1086494 | 2.356323616 | 0.330962733 | 7.119604064 | 1.08E-12 | 2.01E-09 |
| ENSG00000012504.15 | 20.93632437 | 2.933192572 | 0.688058074 | 4.263001459 | 2.02E-05 | 0.002758282 |
| ENSG00000021826.16 | 13367.2312 | 1.782918849 | 0.499680875 | 3.568115048 | 0.000359559 | 0.020870055 |
| ENSG00000037965.6 | 27.40056724 | 2.597604634 | 0.763384276 | 3.402748416 | 0.000667117 | 0.031260939 |
| ENSG00000066279.18 | 376.8948899 | 1.712697988 | 0.449671906 | 3.8087725 | 0.000139658 | 0.010742628 |
| ENSG00000074276.10 | 158.6956445 | 1.58316766 | 0.460104824 | 3.440884719 | 0.000579816 | 0.028437039 |
| ENSG00000077943.8 | 218.3596198 | 2.550537556 | 0.729257781 | 3.497443047 | 0.000469741 | 0.025000524 |
| ENSG00000082556.12 | 90.27568305 | 2.170798805 | 0.444657851 | 4.881953175 | 1.05E-06 | 0.000274505 |
| ENSG00000084453.16 | 157.7459165 | 3.49077367 | 0.74096098 | 4.711143726 | 2.46E-06 | 0.000545783 |
| ENSG00000091583.11 | 2152.189693 | 3.288573352 | 0.665212908 | 4.943640318 | 7.67E-07 | 0.000216229 |
| ENSG00000100181.22 | 141.7242693 | 3.376941415 | 0.586493999 | 5.757844787 | 8.52E-09 | 5.60E-06 |
| ENSG00000107807.13 | 23.78050428 | 2.715067846 | 0.669685806 | 4.054241288 | 5.03E-05 | 0.005001267 |
| ENSG00000109339.23 | 201.3211682 | 1.622318418 | 0.477907932 | 3.394625426 | 0.000687226 | 0.031811437 |
| ENSG00000110900.15 | 21.59918538 | 2.827511411 | 0.665219903 | 4.250491304 | 2.13E-05 | 0.002805129 |
| ENSG00000111181.12 | 21.67017503 | 1.952902339 | 0.570083884 | 3.425640318 | 0.000613352 | 0.029417806 |
| ENSG00000111339.12 | 15.31253871 | 2.467372233 | 0.742903952 | 3.321253343 | 0.000896142 | 0.03756932 |
| ENSG00000112984.12 | 136.6413387 | 2.464386512 | 0.612963624 | 4.020444957 | 5.81E-05 | 0.005619843 |
| ENSG00000113100.10 | 20.80886858 | 2.757357995 | 0.812223078 | 3.394828427 | 0.000686716 | 0.031811437 |
| ENSG00000117724.13 | 374.4885183 | 1.616630128 | 0.436978186 | 3.69956712 | 0.000215968 | 0.014386542 |
| ENSG00000118514.14 | 148.3319298 | 2.495820303 | 0.668307433 | 3.734539195 | 0.000188059 | 0.013128078 |
| ENSG00000122966.16 | 227.5215979 | 1.751250839 | 0.354380885 | 4.941719242 | 7.74E-07 | 0.000216229 |
| ENSG00000124564.17 | 33.0881882 | 2.472639695 | 0.523395968 | 4.724223807 | 2.31E-06 | 0.000519057 |
| ENSG00000130226.17 | 6.520788416 | 5.809317046 | 1.392310872 | 4.17242813 | 3.01E-05 | 0.003590153 |
| ENSG00000130649.10 | 24.97741883 | 2.117472237 | 0.528783558 | 4.004421479 | 6.22E-05 | 0.005949586 |
| ENSG00000131620.17 | 318.5138556 | 1.902217934 | 0.500367218 | 3.801643806 | 0.000143739 | 0.010838664 |
| ENSG00000131849.12 | 55.34913958 | 2.341205225 | 0.386226298 | 6.061744731 | 1.35E-09 | 1.06E-06 |
| ENSG00000132872.12 | 26.27637121 | 3.705773321 | 0.904209137 | 4.098358632 | 4.16E-05 | 0.00448761 |
| ENSG00000132874.14 | 28.53784202 | 1.966319066 | 0.553900351 | 3.549950929 | 0.000385303 | 0.021636407 |
| ENSG00000132967.9 | 17.05369186 | 2.291505134 | 0.697161347 | 3.286907894 | 0.00101294 | 0.040565345 |
| ENSG00000133800.9 | 40.42306975 | 3.277495031 | 0.99029527 | 3.309613939 | 0.000934247 | 0.038472116 |
| ENSG00000135119.14 | 138.6795926 | 2.353382154 | 0.331744764 | 7.093954185 | 1.30E-12 | 2.18E-09 |

| | | | | | | |
|---|---|---|---|---|---|---|
| ENSG00000136011.15 | 25.18619765 | 4.6775075 | 1.240167581 | 3.771673743 | 0.000162156 | 0.011665041 |
| ENSG00000137558.9 | 15.36560727 | 3.448033993 | 0.906732441 | 3.8027028 | 0.000143126 | 0.010838664 |
| ENSG00000138792.10 | 395.6508988 | 1.508744165 | 0.449080604 | 3.359628877 | 0.000780472 | 0.034159866 |
| ENSG00000139209.16 | 196.1187183 | 2.295301577 | 0.593945733 | 3.864497126 | 0.000111318 | 0.008864353 |
| ENSG00000139304.16 | 14.25409904 | 2.843815731 | 0.837960558 | 3.393734592 | 0.000689465 | 0.031811437 |
| ENSG00000139567.12 | 77.86002812 | 1.544155326 | 0.42063987 | 3.670967583 | 0.000241634 | 0.015736317 |
| ENSG00000139973.16 | 157.935978 | 1.620101362 | 0.484815979 | 3.341683099 | 0.000832721 | 0.0354367 |
| ENSG00000142677.4 | 199.9119008 | 1.889088394 | 0.370486117 | 5.098945167 | 3.42E-07 | 0.000118003 |
| ENSG00000147246.10 | 10.61319949 | 3.887433249 | 1.149453967 | 3.381982543 | 0.000719647 | 0.032743947 |
| ENSG00000148386.9 | 16.73036207 | 6.196386497 | 1.515913685 | 4.087558914 | 4.36E-05 | 0.004604207 |
| ENSG00000149124.11 | 137.6270563 | 2.591795213 | 0.443115186 | 5.849032705 | 4.94E-09 | 3.70E-06 |
| ENSG00000150636.17 | 43.71899581 | 1.504481598 | 0.469062219 | 3.207424381 | 0.001339293 | 0.048942961 |
| ENSG00000151491.14 | 269.4629014 | 1.783317842 | 0.490505903 | 3.635670502 | 0.000277259 | 0.017389656 |
| ENSG00000153253.18 | 20.52956541 | 2.988316965 | 0.80357884 | 3.718760146 | 0.000200203 | 0.013603129 |
| ENSG00000155754.15 | 5.396592172 | 5.534037111 | 1.406337659 | 3.935069984 | 8.32E-05 | 0.007321666 |
| ENSG00000156006.5 | 24.6688662 | 3.284588266 | 0.916515991 | 3.583776278 | 0.000338662 | 0.020236291 |
| ENSG00000157152.17 | 201.538287 | 1.619658696 | 0.239419471 | 6.764941413 | 1.33E-11 | 1.94E-08 |
| ENSG00000161031.13 | 130.795591 | 3.592505632 | 0.945434434 | 3.799846403 | 0.000144786 | 0.010846891 |
| ENSG00000161381.14 | 40.17964868 | 1.801646657 | 0.538195365 | 3.347570001 | 0.000815234 | 0.034980242 |
| ENSG00000161798.7 | 24.98529681 | 2.065978034 | 0.503959342 | 4.099493478 | 4.14E-05 | 0.00448761 |
| ENSG00000162692.12 | 145.5770032 | 3.015395762 | 0.597337975 | 5.048056361 | 4.46E-07 | 0.000143951 |
| ENSG00000165169.11 | 110.2434697 | 2.49494185 | 0.549690527 | 4.538811803 | 5.66E-06 | 0.001077725 |
| ENSG00000165899.11 | 38.42455057 | 1.657389735 | 0.473620862 | 3.499401877 | 0.000466303 | 0.024947833 |
| ENSG00000166278.15 | 264.3568962 | 1.54719727 | 0.392175365 | 3.945166903 | 7.97E-05 | 0.007079023 |
| ENSG00000166415.15 | 300.4264757 | 2.03090912 | 0.303440127 | 6.692948415 | 2.19E-11 | 2.97E-08 |
| ENSG00000166863.12 | 153.9825249 | 1.862273271 | 0.54701252 | 3.404443598 | 0.00066299 | 0.031256963 |
| ENSG00000166866.13 | 187.7773921 | 1.775789215 | 0.401415011 | 4.423823636 | 9.70E-06 | 0.001594048 |
| ENSG00000168509.20 | 22.19344236 | 3.172111733 | 0.706022054 | 4.492935758 | 7.02E-06 | 0.001256083 |
| ENSG00000170373.8 | 17.95576664 | 3.289605224 | 1.016160435 | 3.237289222 | 0.00120671 | 0.045891018 |
| ENSG00000171954.13 | 15.18044018 | 3.137465281 | 0.838370429 | 3.742337723 | 0.000182316 | 0.012820387 |
| ENSG00000173535.14 | 95.49748286 | 1.539791839 | 0.377325333 | 4.080806945 | 4.49E-05 | 0.00464688 |
| ENSG00000173578.7 | 13.58916449 | 3.507759956 | 1.007503991 | 3.481633808 | 0.000498365 | 0.025849024 |
| ENSG00000173867.10 | 14.65692597 | 3.072962405 | 0.757807093 | 4.055072106 | 5.01E-05 | 0.005001267 |
| ENSG00000175329.13 | 56.74254818 | 1.51184161 | 0.388423494 | 3.892250682 | 9.93E-05 | 0.008196393 |

| | | | | | | |
|---|---|---|---|---|---|---|
| ENSG00000178187.7 | 19.28102217 | 1.896492189 | 0.548711896 | 3.45626221 | 0.000547722 | 0.027567333 |
| ENSG00000179542.16 | 156.0744471 | 2.84715374 | 0.734380342 | 3.876947105 | 0.000105775 | 0.008522232 |
| ENSG00000180532.10 | 9.01200328 | 5.301961638 | 1.487578628 | 3.56415556 | 0.000365029 | 0.020926996 |
| ENSG00000183114.8 | 26.70964588 | 1.805170025 | 0.562220342 | 3.210787462 | 0.001323718 | 0.048705176 |
| ENSG00000184434.8 | 20.47798975 | 3.25858052 | 0.778399213 | 4.186258753 | 2.84E-05 | 0.003473872 |
| ENSG00000185306.13 | 11.43801523 | 2.936840303 | 0.906597889 | 3.239407833 | 0.001197782 | 0.045722064 |
| ENSG00000185666.14 | 60.66047157 | 2.15382895 | 0.663706674 | 3.245151865 | 0.00117388 | 0.04497814 |
| ENSG00000185736.16 | 41.31009361 | 1.55655562 | 0.452543221 | 3.439573384 | 0.000582632 | 0.028437039 |
| ENSG00000186094.17 | 16.57270089 | 3.62813083 | 0.739330128 | 4.90732177 | 9.23E-07 | 0.000247635 |
| ENSG00000186204.15 | 60.19553614 | 5.590343564 | 0.699742965 | 7.989138644 | 1.36E-15 | 5.54E-12 |
| ENSG00000186529.16 | 1967.120979 | 3.766608479 | 0.614279587 | 6.131749383 | 8.69E-10 | 7.09E-07 |
| ENSG00000187510.9 | 27.38440335 | 3.31112426 | 0.616779174 | 5.368411257 | 7.94E-08 | 3.37E-05 |
| ENSG00000189229.11 | 17.50520897 | 2.819984992 | 0.787020631 | 3.583114446 | 0.000339522 | 0.020236291 |
| ENSG00000196090.12 | 668.2208998 | 5.601563909 | 0.41410316 | 13.52697696 | 1.08E-41 | 2.21E-37 |
| ENSG00000198028.4 | 15.95349727 | 2.817919537 | 0.783587873 | 3.596175533 | 0.00032293 | 0.019649564 |
| ENSG00000198300.14 | 37.86512244 | 8.346677367 | 1.321362332 | 6.316721131 | 2.67E-10 | 3.03E-07 |
| ENSG00000203797.10 | 5.662484607 | 5.599128755 | 1.443496128 | 3.878866487 | 0.000104944 | 0.00848883 |
| ENSG00000203799.13 | 183.441802 | 2.169481812 | 0.584878234 | 3.709287998 | 0.000207843 | 0.013936414 |
| ENSG00000204613.11 | 6.252064175 | 4.739065098 | 1.393107979 | 3.401793091 | 0.000669453 | 0.031298464 |
| ENSG00000204970.9 | 9.351439633 | 6.325305815 | 1.568054288 | 4.033856393 | 5.49E-05 | 0.005377148 |
| ENSG00000206195.11 | 9.0706646 | 5.297425951 | 1.363563359 | 3.884987021 | 0.000102335 | 0.008344016 |
| ENSG00000214429.3 | 6.702235293 | 5.852170123 | 1.585025125 | 3.692162369 | 0.000222355 | 0.014668262 |
| ENSG00000214553.10 | 94.85092642 | 1.908179951 | 0.340463647 | 5.604651093 | 2.09E-08 | 1.22E-05 |
| ENSG00000215808.4 | 45.70782802 | 3.482706009 | 0.810994794 | 4.294362964 | 1.75E-05 | 0.002498704 |
| ENSG00000217330.1 | 53.2883736 | 7.039353328 | 1.274505589 | 5.523203185 | 3.33E-08 | 1.70E-05 |
| ENSG00000224597.10 | 214.2024888 | 1.829395996 | 0.348231438 | 5.253391268 | 1.49E-07 | 6.09E-05 |
| ENSG00000224957.6 | 29.69624505 | 3.877246056 | 0.954551018 | 4.061853145 | 4.87E-05 | 0.004961945 |
| ENSG00000225431.1 | 34.23630058 | 2.240105746 | 0.535235391 | 4.185272092 | 2.85E-05 | 0.003473872 |
| ENSG00000226453.2 | 18.57155866 | 2.64102932 | 0.711839212 | 3.710148691 | 0.000207138 | 0.013934957 |
| ENSG00000226476.5 | 10.99314341 | 3.502865919 | 0.990456098 | 3.536619064 | 0.000405284 | 0.022267667 |
| ENSG00000226674.11 | 35.5047516 | 3.28758106 | 0.504309678 | 6.518972771 | 7.08E-11 | 8.49E-08 |
| ENSG00000227722.2 | 9.348581955 | 5.339408688 | 1.321127323 | 4.041554962 | 5.31E-05 | 0.005228735 |
| ENSG00000229970.3 | 8.042661988 | 5.118089042 | 1.331475449 | 3.843922956 | 0.000121083 | 0.009529562 |
| ENSG00000230453.9 | 30.34394002 | 1.85381807 | 0.498949081 | 3.715445407 | 0.000202846 | 0.013691437 |

| | | | | | | |
|---|---|---|---|---|---|---|
| ENSG00000233098.9 | 87.77521625 | 1.834777343 | 0.391538436 | 4.686072101 | 2.78E-06 | 0.000603926 |
| ENSG00000233508.3 | 9.61175466 | 6.373106809 | 1.405817429 | 4.533381561 | 5.80E-06 | 0.00109558 |
| ENSG00000234985.2 | 13.33022011 | 6.843316959 | 1.516696439 | 4.511988546 | 6.42E-06 | 0.001179383 |
| ENSG00000236532.6 | 11.95214614 | 2.630349319 | 0.797767271 | 3.297138672 | 0.000976753 | 0.039504217 |
| ENSG00000237978.6 | 9.592268949 | 3.350757894 | 1.007954682 | 3.324314033 | 0.000886363 | 0.037329811 |
| ENSG00000239922.2 | 48.33830371 | 1.962989765 | 0.430625494 | 4.558461567 | 5.15E-06 | 0.000990926 |
| ENSG00000240086.7 | 26.96762794 | 2.084427743 | 0.623415491 | 3.343561033 | 0.000827105 | 0.0353453 |
| ENSG00000241644.2 | 1082.754535 | 2.031156249 | 0.531404665 | 3.822240153 | 0.000132245 | 0.010249726 |
| ENSG00000242082.2 | 98.18858467 | 1.503954223 | 0.441817948 | 3.40401342 | 0.000664035 | 0.031256963 |
| ENSG00000242781.2 | 4.495362146 | 5.269526169 | 1.448357123 | 3.638278217 | 0.000274467 | 0.017267692 |
| ENSG00000244306.11 | 21.41709775 | 4.353374909 | 0.850365936 | 5.119413568 | 3.06E-07 | 0.000107714 |
| ENSG00000244694.7 | 37.51727658 | 2.210140554 | 0.511649236 | 4.319640091 | 1.56E-05 | 0.002308471 |
| ENSG00000245146.7 | 21.5668606 | 1.751100548 | 0.492200102 | 3.5577005 | 0.000374116 | 0.021124573 |
| ENSG00000248740.6 | 15.79391246 | 4.612686294 | 1.00350717 | 4.596565358 | 4.30E-06 | 0.000893386 |
| ENSG00000248869.7 | 9.352130134 | 6.327279922 | 1.328936864 | 4.761159158 | 1.92E-06 | 0.000445863 |
| ENSG00000248874.5 | 11.30425006 | 6.601601957 | 1.326425091 | 4.976988147 | 6.46E-07 | 0.000193592 |
| ENSG00000250159.7 | 61.02156887 | 3.586669581 | 0.570605185 | 6.28572904 | 3.26E-10 | 3.17E-07 |
| ENSG00000250546.6 | 18.66440435 | 2.950140426 | 0.69516567 | 4.243794755 | 2.20E-05 | 0.002864537 |
| ENSG00000250616.2 | 34.98637454 | 3.327855761 | 0.529309072 | 6.287169324 | 3.23E-10 | 3.17E-07 |
| ENSG00000250682.6 | 25.20388336 | 1.904271388 | 0.557911429 | 3.413214516 | 0.000642014 | 0.030576659 |
| ENSG00000251095.7 | 156.4955109 | 1.989545833 | 0.561317949 | 3.544418697 | 0.00039348 | 0.021854749 |
| ENSG00000251138.7 | 87.53733281 | 2.039771885 | 0.472681872 | 4.315316507 | 1.59E-05 | 0.002337185 |
| ENSG00000251209.9 | 32.80850788 | 2.635687644 | 0.617536495 | 4.268067825 | 1.97E-05 | 0.002734138 |
| ENSG00000251226.1 | 23.42254209 | 1.642072475 | 0.511976708 | 3.207318712 | 0.001339785 | 0.048942961 |
| ENSG00000253706.5 | 13.58208067 | 4.937925046 | 1.149894647 | 4.294241268 | 1.75E-05 | 0.002498704 |
| ENSG00000254101.7 | 5.169376438 | 5.475516844 | 1.607607061 | 3.406004475 | 0.000659211 | 0.031177162 |
| ENSG00000254245.2 | 29.0719355 | 4.078389471 | 0.742502392 | 5.492762734 | 3.96E-08 | 1.88E-05 |
| ENSG00000254349.6 | 5.523439251 | 4.581163589 | 1.430804753 | 3.201809037 | 0.001365675 | 0.049374435 |
| ENSG00000255282.6 | 7.061552534 | 3.934400066 | 1.223927289 | 3.214570098 | 0.0013064 | 0.048329692 |
| ENSG00000256040.2 | 21.07387462 | 2.273091838 | 0.630018489 | 3.607976395 | 0.000308595 | 0.019061792 |
| ENSG00000257185.2 | 3.984052103 | 5.101919851 | 1.522048937 | 3.352007762 | 0.000802278 | 0.034574267 |
| ENSG00000257869.1 | 5.912242181 | 4.675980644 | 1.458865423 | 3.205217268 | 0.001349606 | 0.049213527 |
| ENSG00000258084.6 | 4.072652597 | 5.139830914 | 1.587015706 | 3.238676779 | 0.001200856 | 0.045753727 |
| ENSG00000259075.6 | 7.385412882 | 5.982620153 | 1.606991555 | 3.722869692 | 0.000196971 | 0.013455834 |

| | | | | | | |
|---|---|---|---|---|---|---|
| ENSG00000259129.6 | 47.94418049 | 3.780888456 | 0.79097413 | 4.780040604 | 1.75E-06 | 0.000415406 |
| ENSG00000260930.2 | 3.410204113 | 4.872780187 | 1.508168403 | 3.230925789 | 0.0012339 | 0.046346203 |
| ENSG00000262352.2 | 4.042633981 | 5.126551905 | 1.499430019 | 3.419000447 | 0.000628516 | 0.030003922 |
| ENSG00000262576.3 | 12.41340096 | 3.7381537 | 0.96324033 | 3.880811034 | 0.000104109 | 0.008454789 |
| ENSG00000262943.7 | 66.77385651 | 2.234003931 | 0.57251485 | 3.902089061 | 9.54E-05 | 0.008012578 |
| ENSG00000263711.6 | 15.84677798 | 7.092755467 | 1.282654519 | 5.529747382 | 3.21E-08 | 1.68E-05 |
| ENSG00000263745.7 | 3.77381424 | 5.020824079 | 1.477046914 | 3.399231286 | 0.000675755 | 0.031520822 |
| ENSG00000267313.7 | 91.99826001 | 2.089513205 | 0.433346831 | 4.821803361 | 1.42E-06 | 0.000349392 |
| ENSG00000267327.2 | 55.11746839 | 3.887291601 | 0.531184325 | 7.318159475 | 2.51E-13 | 5.69E-10 |
| ENSG00000268658.5 | 88.36272434 | 3.833566329 | 0.468896313 | 8.175722904 | 2.94E-16 | 1.50E-12 |
| ENSG00000271369.1 | 13.69401803 | 2.681301995 | 0.686593068 | 3.905227304 | 9.41E-05 | 0.007988468 |
| ENSG00000276476.3 | 72.36970937 | 1.654661549 | 0.381599212 | 4.33612412 | 1.45E-05 | 0.002205991 |
| ENSG00000276975.3 | 29.64938448 | 2.890980892 | 0.724652693 | 3.989470984 | 6.62E-05 | 0.006183704 |
| ENSG00000279511.1 | 5.159094591 | 4.453566839 | 1.387621196 | 3.209497559 | 0.001329672 | 0.048836088 |
| ENSG00000281344.1 | 306.630994 | 2.400004954 | 0.614687633 | 3.904430194 | 9.44E-05 | 0.007988468 |
| ENSG00000283633.1 | 39.50730018 | 3.149844278 | 0.903362255 | 3.486800848 | 0.000488835 | 0.025681488 |
| ENSG00000285876.1 | 6.352988119 | 5.773185509 | 1.390075873 | 4.15314417 | 3.28E-05 | 0.003798118 |
| ENSG00000286329.1 | 26.29101883 | 6.848224936 | 1.248868002 | 5.48354584 | 4.17E-08 | 1.93E-05 |
| ENSG00000286339.1 | 135.9737397 | 7.351382623 | 0.897825588 | 8.187985192 | 2.66E-16 | 1.50E-12 |
| ENSG00000287248.1 | 20.51751643 | 5.563854157 | 1.115033041 | 4.989855862 | 6.04E-07 | 0.000183834 |

| Down-regulted genes | | | | | | |
|---|---|---|---|---|---|---|
| **Gene** | **BaseMean** | **log2FoldChange** | **lfcSE** | **stat** | **pvalue** | **padj** |
| ENSG00000004799.8 | 157.66943 | -3.238477961 | 0.706369758 | -4.584678105 | 4.55E-06 | 0.000926833 |
| ENSG00000004948.15 | 19.90643047 | -4.376504698 | 1.070058972 | -4.089965892 | 4.31E-05 | 0.004604207 |
| ENSG00000005961.19 | 120.9677781 | -3.706929492 | 0.520376007 | -7.123559577 | 1.05E-12 | 2.01E-09 |
| ENSG00000005981.13 | 81.13974438 | -3.79148972 | 0.738998815 | -5.130576182 | 2.89E-07 | 0.000107714 |
| ENSG00000008516.17 | 15.29629468 | -4.592208178 | 1.101268407 | -4.169926378 | 3.05E-05 | 0.003590153 |
| ENSG00000011422.12 | 38.62396938 | -2.893042931 | 0.657042449 | -4.403129409 | 1.07E-05 | 0.001686033 |
| ENSG00000012779.11 | 34.23300062 | -2.839123882 | 0.729336426 | -3.892749327 | 9.91E-05 | 0.008196393 |
| ENSG00000018280.17 | 97.10447862 | -4.119115488 | 0.795667988 | -5.1769275 | 2.26E-07 | 8.68E-05 |
| ENSG00000022556.16 | 251.8822545 | -3.585415702 | 0.386766837 | -9.270225264 | 1.86E-20 | 1.89E-16 |
| ENSG00000025708.14 | 120.144074 | -2.570415405 | 0.563738557 | -4.559587729 | 5.13E-06 | 0.000990926 |
| ENSG00000029534.20 | 120.7097269 | -2.011824217 | 0.477443931 | -4.213739223 | 2.51E-05 | 0.003180123 |

| | | | | | | |
|---|---|---|---|---|---|---|
| ENSG00000047457.14 | 13.01370452 | -3.541765415 | 1.072121397 | -3.303511548 | 0.00095482 | 0.039004125 |
| ENSG00000053918.17 | 53.9905912 | -1.60102771 | 0.40822443 | -3.921930174 | 8.78E-05 | 0.007602954 |
| ENSG00000064201.15 | 26.54337729 | -3.366209337 | 0.85553791 | -3.934611544 | 8.33E-05 | 0.007321666 |
| ENSG00000065911.12 | 742.4675734 | -3.055835865 | 0.602385333 | -5.072892212 | 3.92E-07 | 0.000133112 |
| ENSG00000065989.16 | 22.98544926 | -1.842153993 | 0.466546905 | -3.948486154 | 7.86E-05 | 0.007031315 |
| ENSG00000070182.21 | 33.86677719 | -2.224736671 | 0.626524071 | -3.550919706 | 0.000383888 | 0.021616473 |
| ENSG00000070190.13 | 10.46732237 | -3.582935725 | 1.089034715 | -3.290010572 | 0.001001836 | 0.040278952 |
| ENSG00000070669.17 | 856.9137776 | -3.552522339 | 0.777142183 | -4.571264329 | 4.85E-06 | 0.00096882 |
| ENSG00000074370.18 | 97.21439369 | -2.345892136 | 0.462778944 | -5.069141897 | 4.00E-07 | 0.000133536 |
| ENSG00000074416.14 | 34.58465163 | -2.263305245 | 0.606019851 | -3.734704798 | 0.000187936 | 0.013128078 |
| ENSG00000075275.16 | 177.0974065 | -2.120984099 | 0.582277622 | -3.642565021 | 0.000269935 | 0.017144916 |
| ENSG00000077454.16 | 462.0509173 | -1.721699103 | 0.469942634 | -3.663636749 | 0.000248659 | 0.016004153 |
| ENSG00000078081.8 | 41.20918085 | -2.62825073 | 0.77223996 | -3.403411977 | 0.000665499 | 0.031256963 |
| ENSG00000084636.18 | 404.8092359 | -3.537812718 | 0.845861658 | -4.182495665 | 2.88E-05 | 0.003477661 |
| ENSG00000085514.16 | 19.9938559 | -3.343406983 | 0.822034595 | -4.067233912 | 4.76E-05 | 0.004873157 |
| ENSG00000087074.8 | 723.6439961 | -1.537338491 | 0.480446827 | -3.199809852 | 0.001375183 | 0.049526018 |
| ENSG00000087077.14 | 180.2701547 | -1.647072693 | 0.496155884 | -3.319667765 | 0.000901246 | 0.03756932 |
| ENSG00000087266.17 | 1544.195043 | -2.165935014 | 0.623917637 | -3.471507911 | 0.000517544 | 0.026506579 |
| ENSG00000088827.12 | 45.61876296 | -7.264103743 | 2.161713281 | -3.360345615 | 0.00077845 | 0.034159866 |
| ENSG00000089820.15 | 99.22487945 | -2.108248987 | 0.577539717 | -3.650396547 | 0.000261836 | 0.016731223 |
| ENSG00000091592.16 | 54.21150532 | -2.498699289 | 0.669946936 | -3.729697315 | 0.00019171 | 0.013291888 |
| ENSG00000094755.17 | 118.835061 | -5.325044795 | 1.02560796 | -5.192086065 | 2.08E-07 | 8.15E-05 |
| ENSG00000099251.14 | 26.89751642 | -2.761211324 | 0.73966186 | -3.733072464 | 0.000189158 | 0.013159726 |
| ENSG00000100055.21 | 33.58334658 | -4.143480883 | 1.157447592 | -3.579843193 | 0.0003438 | 0.020372176 |
| ENSG00000100351.17 | 20.35826616 | -3.257636987 | 0.729741392 | -4.464097858 | 8.04E-06 | 0.001392841 |
| ENSG00000100368.14 | 32.03235575 | -2.232713542 | 0.667074589 | -3.347022324 | 0.000816846 | 0.034980242 |
| ENSG00000100385.14 | 46.38103785 | -2.093126081 | 0.527032134 | -3.97153408 | 7.14E-05 | 0.006527568 |
| ENSG00000101162.3 | 52.92498754 | -2.810761216 | 0.712527759 | -3.944774335 | 7.99E-05 | 0.007079023 |
| ENSG00000101255.11 | 761.9099034 | -3.128001142 | 0.770771651 | -4.058272172 | 4.94E-05 | 0.004972341 |
| ENSG00000101307.15 | 20.27443178 | -2.923270702 | 0.732150288 | -3.992719458 | 6.53E-05 | 0.006164257 |
| ENSG00000101670.12 | 1556.668884 | -1.506880484 | 0.332651171 | -4.529911859 | 5.90E-06 | 0.001103509 |
| ENSG00000102007.11 | 56.43865677 | -2.223976887 | 0.638639253 | -3.48236798 | 0.000497 | 0.025844011 |
| ENSG00000103313.12 | 23.45362265 | -4.827170501 | 1.144054083 | -4.219355163 | 2.45E-05 | 0.003121326 |
| ENSG00000103460.17 | 63.36725448 | -2.090060803 | 0.650476735 | -3.213121531 | 0.001313007 | 0.048398442 |

| | | | | | | |
|---|---|---|---|---|---|---|
| ENSG00000103569.10 | 12.00361878 | -4.513744027 | 1.024408431 | -4.406195705 | 1.05E-05 | 0.001675341 |
| ENSG00000105610.5 | 11.65076336 | -4.451025903 | 0.912354808 | -4.878612863 | 1.07E-06 | 0.00027566 |
| ENSG00000105641.4 | 318.7459503 | -3.070894454 | 0.80179139 | -3.830041694 | 0.000128122 | 0.009968052 |
| ENSG00000105835.12 | 496.8414967 | -1.954654846 | 0.275888749 | -7.084938594 | 1.39E-12 | 2.18E-09 |
| ENSG00000105963.15 | 28.99262026 | -1.881317385 | 0.51243125 | -3.671355692 | 0.000241267 | 0.015736317 |
| ENSG00000106366.8 | 136.8850369 | -2.72841087 | 0.662374287 | -4.119137661 | 3.80E-05 | 0.004212984 |
| ENSG00000106479.11 | 94.4594435 | -1.665551721 | 0.476215947 | -3.497471538 | 0.000469691 | 0.025000524 |
| ENSG00000106615.10 | 228.1456038 | -1.55713029 | 0.445805671 | -3.492845405 | 0.000477903 | 0.025237245 |
| ENSG00000106617.14 | 522.4932478 | -1.695401552 | 0.434528758 | -3.901701602 | 9.55E-05 | 0.008012578 |
| ENSG00000106948.16 | 1031.120559 | -1.83415427 | 0.545314074 | -3.363482363 | 0.000769658 | 0.03407203 |
| ENSG00000106952.7 | 9.285984706 | -5.498594723 | 1.403443259 | -3.917931621 | 8.93E-05 | 0.007649313 |
| ENSG00000107317.13 | 34.31511758 | -2.247628667 | 0.691051084 | -3.25247832 | 0.001144033 | 0.044250424 |
| ENSG00000108405.4 | 30.6459644 | -4.245477247 | 0.674665339 | -6.292715815 | 3.12E-10 | 3.17E-07 |
| ENSG00000108556.10 | 25.74014606 | -3.274346596 | 0.800314028 | -4.091327254 | 4.29E-05 | 0.004601542 |
| ENSG00000108839.12 | 5.875736397 | -6.348118581 | 1.463880633 | -4.336500148 | 1.45E-05 | 0.002205991 |
| ENSG00000108932.12 | 22.73439254 | -4.371498131 | 1.056185765 | -4.138948163 | 3.49E-05 | 0.0039077 |
| ENSG00000109321.11 | 58.39490437 | -3.000030006 | 0.676868723 | -4.432218398 | 9.33E-06 | 0.001558348 |
| ENSG00000110090.13 | 593.8083458 | -1.667693593 | 0.400937609 | -4.159484063 | 3.19E-05 | 0.003736683 |
| ENSG00000110243.11 | 238.6903354 | -1.783129787 | 0.412050747 | -4.327451897 | 1.51E-05 | 0.002260889 |
| ENSG00000110324.10 | 125.1200506 | -3.101628614 | 0.877488962 | -3.534663964 | 0.000408294 | 0.022372757 |
| ENSG00000110665.11 | 26.61071063 | -3.514884284 | 0.855623462 | -4.107980251 | 3.99E-05 | 0.004374168 |
| ENSG00000111371.16 | 591.2955483 | -1.846535872 | 0.388835103 | -4.748891903 | 2.05E-06 | 0.000468452 |
| ENSG00000111674.9 | 57.3893836 | -1.979270402 | 0.55920805 | -3.539416864 | 0.000401012 | 0.022130054 |
| ENSG00000113739.10 | 525.4063157 | -4.036027471 | 0.612092828 | -6.593815977 | 4.29E-11 | 5.46E-08 |
| ENSG00000113763.12 | 10.62860153 | -2.405011231 | 0.722382738 | -3.329275609 | 0.000870722 | 0.03674699 |
| ENSG00000114013.16 | 13.92921282 | -3.363743259 | 0.997859831 | -3.37095768 | 0.000749074 | 0.03370024 |
| ENSG00000115155.17 | 27.07986556 | -4.129190497 | 1.158746354 | -3.563498157 | 0.000365945 | 0.020926996 |
| ENSG00000115165.10 | 47.2239909 | -3.778924476 | 0.910371745 | -4.150968542 | 3.31E-05 | 0.003802501 |
| ENSG00000115956.10 | 82.64201068 | -3.864372139 | 0.838475727 | -4.608806211 | 4.05E-06 | 0.000851058 |
| ENSG00000116741.8 | 109.3874778 | -1.899760123 | 0.56748371 | -3.347691023 | 0.000814878 | 0.034980242 |
| ENSG00000117707.16 | 204.5926819 | -1.561894529 | 0.446821185 | -3.495569553 | 0.000473051 | 0.025111132 |
| ENSG00000118523.6 | 163.5014726 | -2.775129949 | 0.568049778 | -4.885364019 | 1.03E-06 | 0.000273299 |
| ENSG00000119681.12 | 33.56695875 | -1.516249359 | 0.46418997 | -3.266441453 | 0.001089083 | 0.043190414 |
| ENSG00000120055.7 | 68.77568011 | -2.502155891 | 0.767857275 | -3.258621064 | 0.001119551 | 0.043894178 |

| | | | | | | |
|---|---|---|---|---|---|---|
| ENSG00000120129.6 | 672.6103099 | -2.170016646 | 0.423421196 | -5.124959891 | 2.98E-07 | 0.000107714 |
| ENSG00000120875.9 | 68.74654616 | -1.933532105 | 0.543317982 | -3.558748597 | 0.000372626 | 0.021124573 |
| ENSG00000120907.18 | 16.84844849 | -3.008878691 | 0.835072715 | -3.60313376 | 0.000314404 | 0.019361944 |
| ENSG00000121281.12 | 273.5126671 | -1.533215419 | 0.384389622 | -3.988701393 | 6.64E-05 | 0.006183704 |
| ENSG00000121410.12 | 17.4125629 | -2.2747891 | 0.588785471 | -3.863527909 | 0.000111761 | 0.008864353 |
| ENSG00000121807.5 | 6.333873394 | -6.45922908 | 1.628987305 | -3.965180734 | 7.33E-05 | 0.006614911 |
| ENSG00000122756.15 | 34.8557423 | -2.464751696 | 0.628652869 | -3.920687898 | 8.83E-05 | 0.007602954 |
| ENSG00000123405.14 | 25.2305656 | -1.752345974 | 0.538095852 | -3.256568448 | 0.001127677 | 0.043894178 |
| ENSG00000124491.15 | 34.61864998 | -3.766320136 | 0.683600886 | -5.509530798 | 3.60E-08 | 1.79E-05 |
| ENSG00000124731.13 | 43.84997746 | -4.79353149 | 1.087717567 | -4.406963382 | 1.05E-05 | 0.001675341 |
| ENSG00000125740.14 | 145.3521987 | -2.976070033 | 0.872640257 | -3.410420286 | 0.000648628 | 0.030819678 |
| ENSG00000125968.9 | 250.6694569 | -1.912755278 | 0.553406343 | -3.456330598 | 0.000547583 | 0.027567333 |
| ENSG00000127824.14 | 31.73823349 | -2.254845646 | 0.659130811 | -3.420938012 | 0.000624056 | 0.029860911 |
| ENSG00000128965.13 | 342.6254435 | -4.802749863 | 0.860751807 | -5.579715106 | 2.41E-08 | 1.33E-05 |
| ENSG00000129911.9 | 231.6938527 | -1.819590076 | 0.562220488 | -3.236435021 | 0.001210328 | 0.045916221 |
| ENSG00000129993.15 | 23.8565499 | -2.572053329 | 0.630140937 | -4.081711214 | 4.47E-05 | 0.00464688 |
| ENSG00000130294.16 | 298.8561561 | -3.439693107 | 0.692524585 | -4.966889527 | 6.80E-07 | 0.000200014 |
| ENSG00000130429.15 | 393.5713247 | -2.007619098 | 0.401134501 | -5.004852722 | 5.59E-07 | 0.000172661 |
| ENSG00000130487.9 | 49.51569977 | -4.843036085 | 0.926637684 | -5.226461399 | 1.73E-07 | 6.91E-05 |
| ENSG00000131042.14 | 46.06365756 | -5.07129698 | 1.169143457 | -4.337617381 | 1.44E-05 | 0.002205991 |
| ENSG00000132334.16 | 96.63225715 | -2.090504055 | 0.607821572 | -3.439338371 | 0.000583138 | 0.028437039 |
| ENSG00000133612.19 | 228.3895986 | -1.625037834 | 0.482236212 | -3.369796368 | 0.000752238 | 0.03370024 |
| ENSG00000134258.17 | 36.23685282 | -4.426395237 | 1.101370029 | -4.018990096 | 5.84E-05 | 0.005619843 |
| ENSG00000134516.17 | 77.91187698 | -3.016831449 | 0.904711687 | -3.334577736 | 0.00085429 | 0.036128322 |
| ENSG00000135185.12 | 128.2446793 | -2.031436686 | 0.352119087 | -5.76917514 | 7.97E-09 | 5.41E-06 |
| ENSG00000135269.18 | 377.2194157 | -2.616778633 | 0.542403295 | -4.824415072 | 1.40E-06 | 0.00034905 |
| ENSG00000135604.10 | 38.31781284 | -2.563344756 | 0.707570547 | -3.622740894 | 0.000291498 | 0.018061913 |
| ENSG00000135842.17 | 604.5889739 | -2.336970408 | 0.670618342 | -3.484799416 | 0.000492506 | 0.025741652 |
| ENSG00000136327.7 | 42.14569189 | -2.547051955 | 0.642002263 | -3.96735666 | 7.27E-05 | 0.006592469 |
| ENSG00000136826.15 | 95.61319477 | -2.548618508 | 0.573852041 | -4.441246746 | 8.94E-06 | 0.001519273 |
| ENSG00000136997.19 | 67.02470769 | -1.802385925 | 0.477485585 | -3.774744165 | 0.000160172 | 0.01161655 |
| ENSG00000137285.10 | 184.8687983 | -2.118568569 | 0.439102377 | -4.824771356 | 1.40E-06 | 0.00034905 |
| ENSG00000137331.12 | 76.35591617 | -1.884907752 | 0.560240956 | -3.364459047 | 0.000766939 | 0.03407203 |
| ENSG00000137801.11 | 1609.581844 | -3.349277885 | 0.724496427 | -4.622904628 | 3.78E-06 | 0.000803477 |

| | | | | | | |
|---|---|---|---|---|---|---|
| ENSG00000137959.16 | 104.5769459 | -4.99687681 | 1.057856595 | -4.723586195 | 2.32E-06 | 0.000519057 |
| ENSG00000137965.11 | 31.34452874 | -4.69079975 | 1.279587731 | -3.665868025 | 0.000246501 | 0.015951356 |
| ENSG00000139269.3 | 108.4357643 | -2.528652671 | 0.530393342 | -4.767504548 | 1.87E-06 | 0.000437018 |
| ENSG00000139514.13 | 621.8365975 | -2.465511993 | 0.539287457 | -4.571795544 | 4.84E-06 | 0.00096882 |
| ENSG00000140379.8 | 12.42392634 | -4.041381883 | 1.033053207 | -3.912075249 | 9.15E-05 | 0.007804461 |
| ENSG00000140678.16 | 83.13413532 | -4.465162322 | 1.077355427 | -4.14455825 | 3.40E-05 | 0.003855627 |
| ENSG00000141506.13 | 60.06123852 | -3.192441737 | 0.923276469 | -3.457731076 | 0.000544745 | 0.027553546 |
| ENSG00000141582.15 | 339.9341965 | -2.628147412 | 0.619419502 | -4.24292003 | 2.21E-05 | 0.002864537 |
| ENSG00000141682.11 | 52.03571743 | -1.853498334 | 0.550334062 | -3.367951326 | 0.00075729 | 0.033852176 |
| ENSG00000141968.8 | 17.37041209 | -2.840830631 | 0.79678856 | -3.565350677 | 0.00036337 | 0.020926996 |
| ENSG00000142102.16 | 122.718893 | -2.247142116 | 0.597468225 | -3.761107325 | 0.000169163 | 0.012099 |
| ENSG00000142185.16 | 27.97122645 | -3.5276108 | 0.798271399 | -4.419061993 | 9.91E-06 | 0.001616536 |
| ENSG00000142552.8 | 204.7267853 | -1.957448144 | 0.537625813 | -3.64091176 | 0.000271674 | 0.017144916 |
| ENSG00000143632.14 | 11.91585656 | -3.130916203 | 0.87870725 | -3.563093629 | 0.00036651 | 0.020926996 |
| ENSG00000145423.5 | 36.37496932 | -3.530813387 | 1.094833729 | -3.224976811 | 0.001259829 | 0.04686195 |
| ENSG00000146112.12 | 387.5877873 | -1.5688424 | 0.47380354 | -3.311166482 | 0.000929079 | 0.038372637 |
| ENSG00000146592.17 | 23.70176136 | -1.843562732 | 0.485016925 | -3.801027628 | 0.000144097 | 0.010838664 |
| ENSG00000146678.10 | 9.171392425 | -5.474414561 | 1.617027363 | -3.385480473 | 0.000710538 | 0.032484926 |
| ENSG00000147872.10 | 1695.173847 | -4.052433908 | 0.803152963 | -5.045656424 | 4.52E-07 | 0.000143951 |
| ENSG00000148339.12 | 508.094609 | -1.870405243 | 0.567061881 | -3.2984147 | 0.000972324 | 0.039430443 |
| ENSG00000148408.13 | 65.68158001 | -1.752002292 | 0.45984832 | -3.809956924 | 0.000138991 | 0.01073179 |
| ENSG00000148488.17 | 19.76836002 | -4.086656439 | 1.137510842 | -3.59263076 | 0.000327356 | 0.019800685 |
| ENSG00000148908.15 | 40.20376055 | -2.379948084 | 0.423919459 | -5.614151539 | 1.98E-08 | 1.18E-05 |
| ENSG00000149781.12 | 73.96998725 | -2.542588621 | 0.75601248 | -3.363156943 | 0.000770565 | 0.03407203 |
| ENSG00000150594.7 | 17.60253204 | -2.569270046 | 0.637815434 | -4.02823436 | 5.62E-05 | 0.005480985 |
| ENSG00000150681.10 | 14.4774392 | -4.225026605 | 1.134549114 | -3.723969772 | 0.000196114 | 0.013455834 |
| ENSG00000151650.8 | 3.386858539 | -5.535659895 | 1.67837734 | -3.298221302 | 0.000972994 | 0.039430443 |
| ENSG00000151952.16 | 12.32466143 | -2.218589862 | 0.612409962 | -3.622720071 | 0.000291521 | 0.018061913 |
| ENSG00000153404.14 | 31.22843419 | -1.989873118 | 0.57798014 | -3.442805346 | 0.000575714 | 0.028437039 |
| ENSG00000154118.13 | 70.6234753 | -1.544781639 | 0.378671653 | -4.079475253 | 4.51E-05 | 0.00464688 |
| ENSG00000154451.14 | 26.06739748 | -4.249932622 | 1.074668965 | -3.954643486 | 7.66E-05 | 0.006882865 |
| ENSG00000155307.18 | 17.62737367 | -3.528380834 | 1.025838786 | -3.439508119 | 0.000582772 | 0.028437039 |
| ENSG00000155897.10 | 9.924189264 | -3.668045556 | 1.046261185 | -3.505860303 | 0.000455134 | 0.024478765 |
| ENSG00000158764.7 | 97.87916906 | -1.950156302 | 0.427382506 | -4.563023227 | 5.04E-06 | 0.000988276 |

| | | | | | | |
|---|---|---|---|---|---|---|
| ENSG00000159399.9 | 1318.570516 | -2.094073321 | 0.476812813 | -4.391814282 | 1.12E-05 | 0.001762568 |
| ENSG00000159784.17 | 14.01362385 | -2.815497949 | 0.673693843 | -4.179195012 | 2.93E-05 | 0.00350776 |
| ENSG00000159840.16 | 665.6208847 | -1.588408257 | 0.43358827 | -3.663402277 | 0.000248887 | 0.016004153 |
| ENSG00000160352.16 | 256.3641983 | -2.899298383 | 0.464983068 | -6.235277331 | 4.51E-10 | 4.18E-07 |
| ENSG00000160360.13 | 305.5771751 | -2.495065748 | 0.771665175 | -3.23335279 | 0.001223464 | 0.046098123 |
| ENSG00000160469.17 | 188.2701119 | -2.258422336 | 0.680233532 | -3.320069108 | 0.000899952 | 0.03756932 |
| ENSG00000160813.7 | 79.17472544 | -2.128468351 | 0.651488915 | -3.267082987 | 0.001086618 | 0.043176659 |
| ENSG00000160991.16 | 93.94165429 | -1.859536954 | 0.300437707 | -6.189425995 | 6.04E-10 | 5.35E-07 |
| ENSG00000162367.11 | 55.65908407 | -1.649874515 | 0.40732698 | -4.050491608 | 5.11E-05 | 0.005057423 |
| ENSG00000162511.8 | 290.2502032 | -2.656230401 | 0.744443002 | -3.5680776 | 0.00035961 | 0.020870055 |
| ENSG00000162711.17 | 40.55284844 | -4.364779665 | 1.23042004 | -3.547389933 | 0.000389068 | 0.021787265 |
| ENSG00000162804.14 | 14.46840973 | -2.044287247 | 0.61692018 | -3.31369813 | 0.000920709 | 0.038145777 |
| ENSG00000163734.4 | 6.469900097 | -3.271751066 | 0.972270344 | -3.365063108 | 0.000765262 | 0.03407203 |
| ENSG00000163736.4 | 19.65740612 | -4.327283492 | 0.947284063 | -4.568094893 | 4.92E-06 | 0.000974033 |
| ENSG00000163737.4 | 43.83968582 | -2.71341248 | 0.777219594 | -3.491178685 | 0.000480895 | 0.025329596 |
| ENSG00000164406.8 | 803.3735542 | -1.863814016 | 0.529488807 | -3.520025338 | 0.000431506 | 0.023455421 |
| ENSG00000164707.15 | 7.29871653 | -3.792165737 | 1.147290075 | -3.305324275 | 0.000948666 | 0.038908656 |
| ENSG00000164897.14 | 233.7962487 | -1.742529934 | 0.501636714 | -3.473688994 | 0.000513356 | 0.026424854 |
| ENSG00000165125.20 | 86.19697368 | -2.350752574 | 0.479026187 | -4.907357134 | 9.23E-07 | 0.000247635 |
| ENSG00000165702.14 | 15.95949732 | -2.402296421 | 0.701200753 | -3.425975242 | 0.000612596 | 0.029417806 |
| ENSG00000165802.22 | 263.3043961 | -1.776829936 | 0.497689396 | -3.570158314 | 0.000356766 | 0.020870055 |
| ENSG00000166091.21 | 4.54742399 | -3.821448046 | 1.173520706 | -3.256395925 | 0.001128363 | 0.043894178 |
| ENSG00000166501.14 | 82.24444717 | -2.326060545 | 0.629531186 | -3.69490916 | 0.000219965 | 0.014557711 |
| ENSG00000167207.13 | 19.26843348 | -1.921684803 | 0.552571122 | -3.47771486 | 0.000505708 | 0.026097071 |
| ENSG00000167208.15 | 12.71994216 | -4.330112575 | 1.202414987 | -3.60117981 | 0.000316776 | 0.0193909 |
| ENSG00000167772.12 | 252.4208021 | -1.582221707 | 0.487356478 | -3.246538785 | 0.001168175 | 0.044928463 |
| ENSG00000167851.15 | 19.79927692 | -2.990647855 | 0.88288769 | -3.387348005 | 0.000705718 | 0.032399454 |
| ENSG00000168209.5 | 2307.375545 | -3.42183175 | 0.589412614 | -5.805494603 | 6.42E-09 | 4.51E-06 |
| ENSG00000168229.4 | 3.372146001 | -4.752540564 | 1.479085639 | -3.213161185 | 0.001312826 | 0.048398442 |
| ENSG00000168268.11 | 752.6568114 | -1.526659173 | 0.2677594 | -5.701608136 | 1.19E-08 | 7.56E-06 |
| ENSG00000169508.7 | 29.69815441 | -2.779044274 | 0.80362673 | -3.458128222 | 0.000543942 | 0.027553546 |
| ENSG00000169704.4 | 3.527897582 | -5.601699689 | 1.506719478 | -3.717811955 | 0.000200956 | 0.013608911 |
| ENSG00000170345.10 | 463.1077342 | -3.481380684 | 0.828123193 | -4.203940567 | 2.62E-05 | 0.003260295 |
| ENSG00000171466.10 | 112.8983069 | -2.902473734 | 0.370431819 | -7.835379095 | 4.67E-15 | 1.59E-11 |

| | | | | | | |
|---|---|---|---|---|---|---|
| ENSG00000171502.15 | 14.75982863 | -3.399773181 | 0.671843035 | -5.060368279 | 4.18E-07 | 0.000137575 |
| ENSG00000171522.6 | 28.24571447 | -1.613542693 | 0.503015527 | -3.207739337 | 0.001337827 | 0.048942961 |
| ENSG00000171860.5 | 10.93886091 | -3.260915969 | 0.990318938 | -3.29279371 | 0.000991972 | 0.040040323 |
| ENSG00000172059.11 | 134.6763623 | -2.344653739 | 0.475432758 | -4.931620089 | 8.16E-07 | 0.000224638 |
| ENSG00000172159.16 | 26.21982586 | -1.861542909 | 0.491316441 | -3.788887878 | 0.000151323 | 0.011251318 |
| ENSG00000172216.6 | 321.2450408 | -1.665710399 | 0.49582586 | -3.359466567 | 0.000780931 | 0.034159866 |
| ENSG00000172243.17 | 47.75279887 | -4.654060804 | 1.041238601 | -4.469735178 | 7.83E-06 | 0.00137621 |
| ENSG00000172322.14 | 14.36252372 | -4.744050168 | 1.457470128 | -3.254989641 | 0.001133965 | 0.044028082 |
| ENSG00000172354.10 | 841.6650728 | -1.50789814 | 0.438079665 | -3.442063765 | 0.000577294 | 0.028437039 |
| ENSG00000172432.19 | 538.0270519 | -2.113000065 | 0.660277835 | -3.200168102 | 0.001373475 | 0.049526018 |
| ENSG00000173110.8 | 21.82817556 | -2.984654847 | 0.822898244 | -3.627003545 | 0.000286729 | 0.017873674 |
| ENSG00000173237.4 | 278.4388645 | -3.518198691 | 0.820942693 | -4.285559418 | 1.82E-05 | 0.002562479 |
| ENSG00000174672.16 | 90.69252184 | -2.542107915 | 0.65781601 | -3.864466474 | 0.000111332 | 0.008864353 |
| ENSG00000175084.12 | 17.03328951 | -2.548628166 | 0.735349431 | -3.465873579 | 0.000528512 | 0.027000459 |
| ENSG00000175592.9 | 5.766516924 | -3.346529963 | 1.035987064 | -3.230281612 | 0.001236683 | 0.046346203 |
| ENSG00000176845.13 | 121.7209718 | -1.54375616 | 0.371455737 | -4.155962629 | 3.24E-05 | 0.003773029 |
| ENSG00000177989.13 | 16.24250856 | -2.728836056 | 0.66401742 | -4.10958504 | 3.96E-05 | 0.004367363 |
| ENSG00000178562.18 | 10.88242399 | -5.170600358 | 1.248271683 | -4.142207526 | 3.44E-05 | 0.003873848 |
| ENSG00000182648.13 | 44.92399934 | -2.138841777 | 0.384327211 | -5.565158325 | 2.62E-08 | 1.40E-05 |
| ENSG00000183010.17 | 684.2574317 | -5.25876478 | 0.688768693 | -7.635022947 | 2.26E-14 | 5.75E-11 |
| ENSG00000183439.9 | 17.65753736 | -4.880261602 | 0.857871001 | -5.688805887 | 1.28E-08 | 7.90E-06 |
| ENSG00000184261.4 | 101.4132382 | -1.824556053 | 0.564056519 | -3.234704311 | 0.001217688 | 0.045965454 |
| ENSG00000185215.9 | 87.14675461 | -2.06112807 | 0.61429549 | -3.355271368 | 0.000792872 | 0.034241305 |
| ENSG00000185245.8 | 9.938928671 | -4.599995647 | 1.028821878 | -4.471129305 | 7.78E-06 | 0.00137621 |
| ENSG00000185561.10 | 257.1789387 | -1.847371509 | 0.518655978 | -3.561843661 | 0.00036826 | 0.02096817 |
| ENSG00000185862.7 | 48.45842685 | -3.903540773 | 1.208611689 | -3.229772481 | 0.001238888 | 0.046346203 |
| ENSG00000185885.16 | 62.6960419 | -2.363058696 | 0.539848548 | -4.377262296 | 1.20E-05 | 0.001870027 |
| ENSG00000185950.9 | 472.0653353 | -2.144449922 | 0.396299477 | -5.411185342 | 6.26E-08 | 2.83E-05 |
| ENSG00000186407.7 | 54.86680946 | -3.743562959 | 1.08535784 | -3.449150889 | 0.000562352 | 0.028026872 |
| ENSG00000187840.5 | 577.0744147 | -1.926644188 | 0.591603887 | -3.256645585 | 0.001127371 | 0.043894178 |
| ENSG00000188536.13 | 218.542599 | -4.142303822 | 0.864937586 | -4.789136106 | 1.68E-06 | 0.000401687 |
| ENSG00000196411.10 | 3149.853533 | -1.869705822 | 0.467892023 | -3.99601987 | 6.44E-05 | 0.006107267 |
| ENSG00000196453.8 | 132.4269008 | -1.785305537 | 0.419154428 | -4.259302583 | 2.05E-05 | 0.002768251 |
| ENSG00000196517.11 | 302.2898959 | -2.435271598 | 0.698973321 | -3.484069455 | 0.000493851 | 0.025745947 |

| | | | | | | |
|---|---|---|---|---|---|---|
| ENSG00000197405.8 | 94.18097152 | -3.924169079 | 0.870098839 | -4.510026793 | 6.48E-06 | 0.001179714 |
| ENSG00000197632.9 | 8.141548874 | -6.803355625 | 1.631514425 | -4.169963515 | 3.05E-05 | 0.003590153 |
| ENSG00000197993.9 | 14.47812867 | -2.43900869 | 0.570294796 | -4.276750742 | 1.90E-05 | 0.002647699 |
| ENSG00000198650.11 | 441.9556545 | -2.830805273 | 0.660246594 | -4.28749697 | 1.81E-05 | 0.002557877 |
| ENSG00000204054.14 | 206.5771985 | -2.024514783 | 0.498908493 | -4.057887991 | 4.95E-05 | 0.004972341 |
| ENSG00000204099.11 | 248.4882707 | -2.38217348 | 0.582631832 | -4.088642859 | 4.34E-05 | 0.004604207 |
| ENSG00000204389.10 | 862.3668793 | -2.709357972 | 0.765578394 | -3.538968697 | 0.000401693 | 0.022130054 |
| ENSG00000204420.10 | 29.16039546 | -4.274860329 | 0.791793903 | -5.398955856 | 6.70E-08 | 2.91E-05 |
| ENSG00000204424.9 | 5.383508968 | -6.219159679 | 1.354303108 | -4.592147537 | 4.39E-06 | 0.000903295 |
| ENSG00000204482.10 | 10.41842468 | -4.656165104 | 1.338514008 | -3.478607677 | 0.000504026 | 0.026076304 |
| ENSG00000204983.14 | 43.10381029 | -3.788084913 | 0.739609673 | -5.121735219 | 3.03E-07 | 0.000107714 |
| ENSG00000205710.4 | 216.6412692 | -2.299139464 | 0.563435104 | -4.080575469 | 4.49E-05 | 0.00464688 |
| ENSG00000205795.4 | 13.652558 | -2.071909425 | 0.634580359 | -3.265007176 | 0.001094613 | 0.043325421 |
| ENSG00000206172.8 | 6.632532217 | -5.775556146 | 1.407832839 | -4.102444543 | 4.09E-05 | 0.004456224 |
| ENSG00000206190.11 | 11.19586725 | -2.540382676 | 0.752003956 | -3.378150681 | 0.000729751 | 0.033129712 |
| ENSG00000206432.4 | 37.58597079 | -1.726056997 | 0.455649971 | -3.788120502 | 0.000151791 | 0.011251318 |
| ENSG00000215146.5 | 4.781812735 | -6.053162711 | 1.422882976 | -4.254153583 | 2.10E-05 | 0.002795689 |
| ENSG00000215218.4 | 28.28047598 | -2.044260336 | 0.569255532 | -3.591111936 | 0.00032927 | 0.019857524 |
| ENSG00000223756.6 | 6.664209722 | -5.781746174 | 1.336731569 | -4.325285874 | 1.52E-05 | 0.002266555 |
| ENSG00000223802.7 | 65.44886505 | -1.735987503 | 0.534910981 | -3.245376451 | 0.001172955 | 0.04497814 |
| ENSG00000224397.7 | 12.32527531 | -4.600507723 | 1.227010043 | -3.749364358 | 0.000177283 | 0.012547722 |
| ENSG00000225968.7 | 205.6490037 | -1.557915913 | 0.420548289 | -3.704487576 | 0.000211818 | 0.01415642 |
| ENSG00000229644.6 | 35.13397667 | -2.799755312 | 0.52395549 | -5.343498383 | 9.12E-08 | 3.79E-05 |
| ENSG00000233297.4 | 9.800841252 | -3.701931099 | 1.037680049 | -3.567507252 | 0.000360393 | 0.020870055 |
| ENSG00000235568.6 | 28.81836829 | -3.335619218 | 1.034251557 | -3.225152714 | 0.001259055 | 0.04686195 |
| ENSG00000236404.10 | 8.846034963 | -3.823272806 | 0.981961174 | -3.893507103 | 9.88E-05 | 0.008196393 |
| ENSG00000239911.2 | 43.6091832 | -1.96832248 | 0.551606589 | -3.568344756 | 0.000359244 | 0.020870055 |
| ENSG00000241685.10 | 451.4934219 | -1.546644807 | 0.329058547 | -4.70021162 | 2.60E-06 | 0.000569639 |
| ENSG00000248323.7 | 19.79903214 | -5.609600252 | 1.264537558 | -4.43608829 | 9.16E-06 | 0.001543259 |
| ENSG00000249364.6 | 41.80401913 | -1.525379739 | 0.461324899 | -3.306519426 | 0.000944628 | 0.038821164 |
| ENSG00000255197.6 | 12.76246679 | -3.862475755 | 1.14443269 | -3.375013479 | 0.000738121 | 0.033358004 |
| ENSG00000259171.1 | 109.3107567 | -1.603477351 | 0.392473135 | -4.085572253 | 4.40E-05 | 0.004619846 |
| ENSG00000259207.7 | 156.6746632 | -2.214024925 | 0.446649924 | -4.956958022 | 7.16E-07 | 0.000205578 |
| ENSG00000259803.7 | 11.10877206 | -3.479423335 | 1.041056765 | -3.34220328 | 0.000831162 | 0.0354367 |

| | | | | | | |
|---|---|---|---|---|---|---|
| ENSG00000260997.1 | 9.517178978 | -4.961760683 | 1.245323299 | -3.984315308 | 6.77E-05 | 0.006270393 |
| ENSG00000261455.1 | 15.8299009 | -2.372628193 | 0.628187727 | -3.776941335 | 0.000158766 | 0.01161655 |
| ENSG00000262001.1 | 13.71591457 | -2.911445984 | 0.818275755 | -3.558025478 | 0.000373653 | 0.021124573 |
| ENSG00000262227.1 | 21.43179954 | -2.574492327 | 0.718772463 | -3.581790424 | 0.000341248 | 0.020279851 |
| ENSG00000265206.5 | 6.82773486 | -4.411102486 | 1.275051923 | -3.459547337 | 0.000541084 | 0.027553546 |
| ENSG00000267368.1 | 101.3138259 | -3.044279201 | 0.829839387 | -3.668516161 | 0.000243962 | 0.015837347 |
| ENSG00000267519.6 | 191.5851685 | -3.58002196 | 0.580902049 | -6.162866812 | 7.14E-10 | 6.07E-07 |
| ENSG00000267943.1 | 10.23360139 | -3.769349657 | 0.947559429 | -3.977955939 | 6.95E-05 | 0.006411298 |
| ENSG00000272079.2 | 15.70883553 | -2.97373297 | 0.787040816 | -3.778371983 | 0.000157857 | 0.01161655 |
| ENSG00000272398.6 | 1314.837397 | -2.172748387 | 0.558659425 | -3.889218173 | 0.000100568 | 0.008266014 |
| ENSG00000272512.1 | 8.759756264 | -2.357511883 | 0.710166281 | -3.319661811 | 0.000901266 | 0.03756932 |
| ENSG00000274750.2 | 43.77771003 | -6.470695922 | 0.839040573 | -7.712017909 | 1.24E-14 | 3.61E-11 |
| ENSG00000275620.1 | 14.31714038 | -2.901739562 | 0.808423046 | -3.589382535 | 0.000331462 | 0.01993075 |
| ENSG00000275896.5 | 11.34639527 | -6.565185698 | 1.280749371 | -5.126050302 | 2.96E-07 | 0.000107714 |
| ENSG00000276107.1 | 5.083948984 | -5.362369147 | 1.429462717 | -3.751317947 | 0.000175907 | 0.01249372 |
| ENSG00000276805.2 | 53.16804457 | -1.800974449 | 0.495742155 | -3.632885422 | 0.000280269 | 0.017524581 |
| ENSG00000280407.2 | 7.176707138 | -3.212717918 | 0.93081201 | -3.451521773 | 0.000557435 | 0.027918306 |
| ENSG00000284707.2 | 45.14085695 | -2.947412712 | 0.877513315 | -3.358823918 | 0.000782749 | 0.034166083 |
