## Supplementary figures and images for "Human Induced Pluripotent Stem Cell based Hepatic-Modeling of Lipid metabolism associated TM6SF2 E167K variant"

### Supplementary Table 1

**Supplementary table 1: Characteristics of human cells used in this study.**

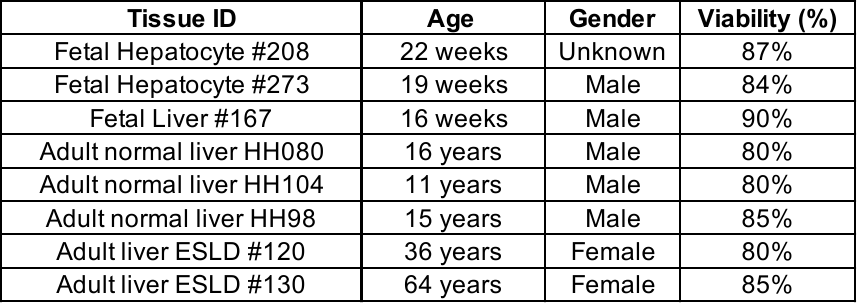
