## Supplementary Table 2 for "Human Induced Pluripotent Stem Cell based Hepatic-Modeling of Lipid metabolism associated TM6SF2 E167K variant"

**Supplementary table 2. Taqman® Gene Expression Assay IDs used for qPCR and genotyping**

| **Target gene** | **Gene Expression Assay ID** | **Company** |
| --- | --- | --- |
| Sox17 | Hs00751752_s1 | Life Technologies |
| Sox9 | Hs01001343_g1 | Life Technologies |
| HNF1ß | Hs01001602_m1 | Life Technologies |
| ITPR3 | Hs01573539_m1 | Life Technologies |
| HNF4a | Hs00604431_m1 | Life Technologies |
| UGT1A1 | Hs02511055_s1 | Life Technologies |
| LXR | Hs00172885_m1 | Life Technologies |
| CFTR | Hs00357011_m1 | Life Technologies |
| LRRC8A | Hs01555916_m1 | Life Technologies |
| TMEM16A | Hs00216121_m1 | Life Technologies |
| ß-actin | Hs01060665_g1 | Life Technologies |
| BSEP | Hs00184824_m1 | Life Technologies |
| MRP2 | Hs00166123_m1 | Life Technologies |
| ACC | Hs01046047_m1 | Life Technologies |
| FASN | Hs01005622_m1 | Life Technologies |
| SCTR | Hs01085380_m1 | Life Technologies |
| AE2 | Hs01586776_m1 | Life Technologies |
| AQP1 | Hs01028916_m1 | Life Technologies |
| CHRM3 | Hs00265216_s1 | Life Technologies |
| P2Y1R | Hs00704965_s1 | Life Technologies |
| c-Myc | Hs00153408_m1 | Life Technologies |
| Sox2 | Hs01053049_s1 | Life Technologies |
| Nanog | Hs02387400_g1 | Life Technologies |
| Oct3/4 | Hs04260367_gH | Life Technologies |
| Lin28A | Hs00702808_s1 | Life Technologies |
| HSP70 | Hs00382884_m1 | Life Technologies |
| FOXA1 | Hs04187555_m1 | Life Technologies |
| ELOVL6 | Hs00907564_m1 | Life Technologies |
| EGFR | Hs01076090_m1 | Life Technologies |
| PPARA | Hs00947536_m1 | Life Technologies |
| FOXA2 | Hs00232764_m1 | Life Technologies |
| Srebf1 | Hs01088691_m1 | Life Technologies |
| RXR | Hs00172885_m1 | Life Technologies |
| TM6SF2 rs58542926 | C__89463510_10 | Life Technologies |
| PNPLA3 rs738409 | C______7241_10 | Life Technologies |
| MBOAT7 rs62641738 | C___8716820_10 | Life Technologies |
| GCKR rs780094 | C___2862873_10 | Life Technologies |
| HSD17B13 rs72613567 | ANWDFW4 | Life Technologies |
| MTARC1 rs2642438 | C___1235772_10 | Life Technologies |
