## Supplementary Table 3 for "Human Induced Pluripotent Stem Cell based Hepatic-Modeling of Lipid metabolism associated TM6SF2 E167K variant"

**Supplementary table 3. Antibodies and dilutions used for Western blot and Immunofluorescence staining.**

| **Target protein** | **Catalog number** | **Company** | **Dilution** | **Host** |
| --- | --- | --- | --- | --- |
| anti-Sox17 | NL1924R | R&D | 1:50 | Goat |
| anti-HNF4α | Ab41898 | Abcam | 1:500 | Mouse |
| anti-Alb | A80-229A | Bethyl | 1:100 | Goat |
| anti-AFP | ZSA06 | Invitrogen | 1:300 | Mouse |
| anti-TRA-1-60 | 560218 | BD Biosciences | 1:10 | Mouse |
| anti-TM6SF2 | PA5-69304 | Invitrogen | 1:100 | Rabbit |
| anti-HSPA5 | NBP1-06274 | Novus | 1:1,000 | Rabbit |
| anti-XBP1 | Ab37152 | Abcam | 1:50 | Rabbit |
| anti-OCT3/4 | SC-9081 | Santa-Cruz | 1:250 | Mouse |
| anti-SSEA4 | 14-8843-80 | Invitrogen | 1:10 | Mouse |
| anti-Nanog | 4893S | Cell signaling | 1:2000 | Rabbit |
| anti-SOX1 | SC022 | R&D | 1:10 | Goat |
| anti-Otx-2 |  |  |  | Goat |
| anti-HAND-1 |  |  |  | Goat |
| anti-Brachyury |  |  |  | Goat |
| anti-GATA-4 |  |  |  | Goat |
| anti-SOX17 |  |  |  | Goat |
| anti-TM6SF2 (WB) | PA5-69304 | Invitrogen | 1:1,500 | Rabbit |
| anti-HSP70 (WB) | Ab2787 | Abcam | 1:5,000 | Mouse |
| anti-ApoB (WB) | SC-393636 | Santa-Cruz | 1:1,000 | Mouse |
| anti-GAPDH (WB) | 60004-1-1g | ProteinTech | 1:10,000 | Mouse |
