## Supplementary Table 5 for "Human Induced Pluripotent Stem Cell based Hepatic-Modeling of Lipid metabolism associated TM6SF2 E167K variant"

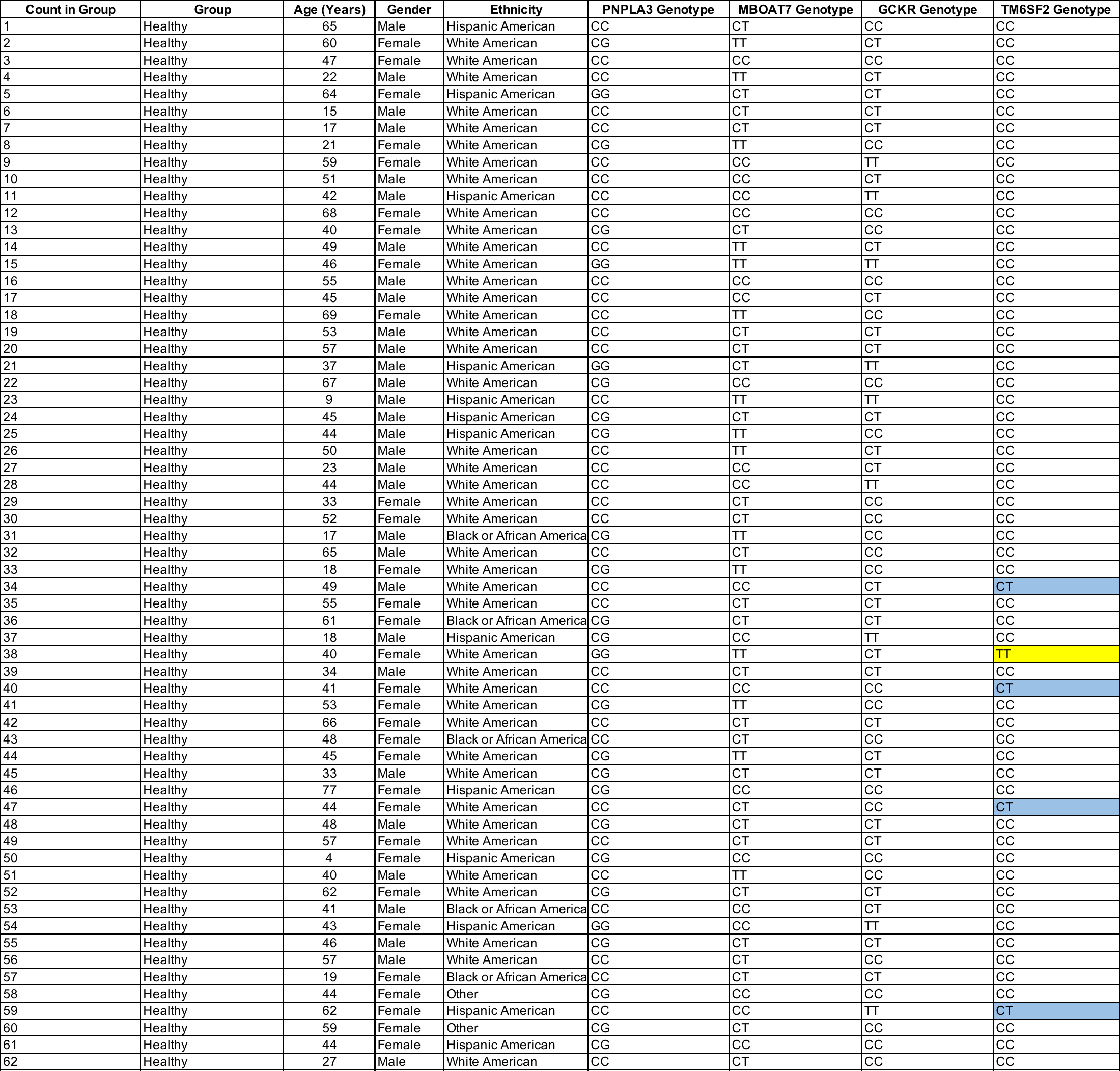
**Supplementary table 5a: Characteristics of healthy cohorts analyzed.**

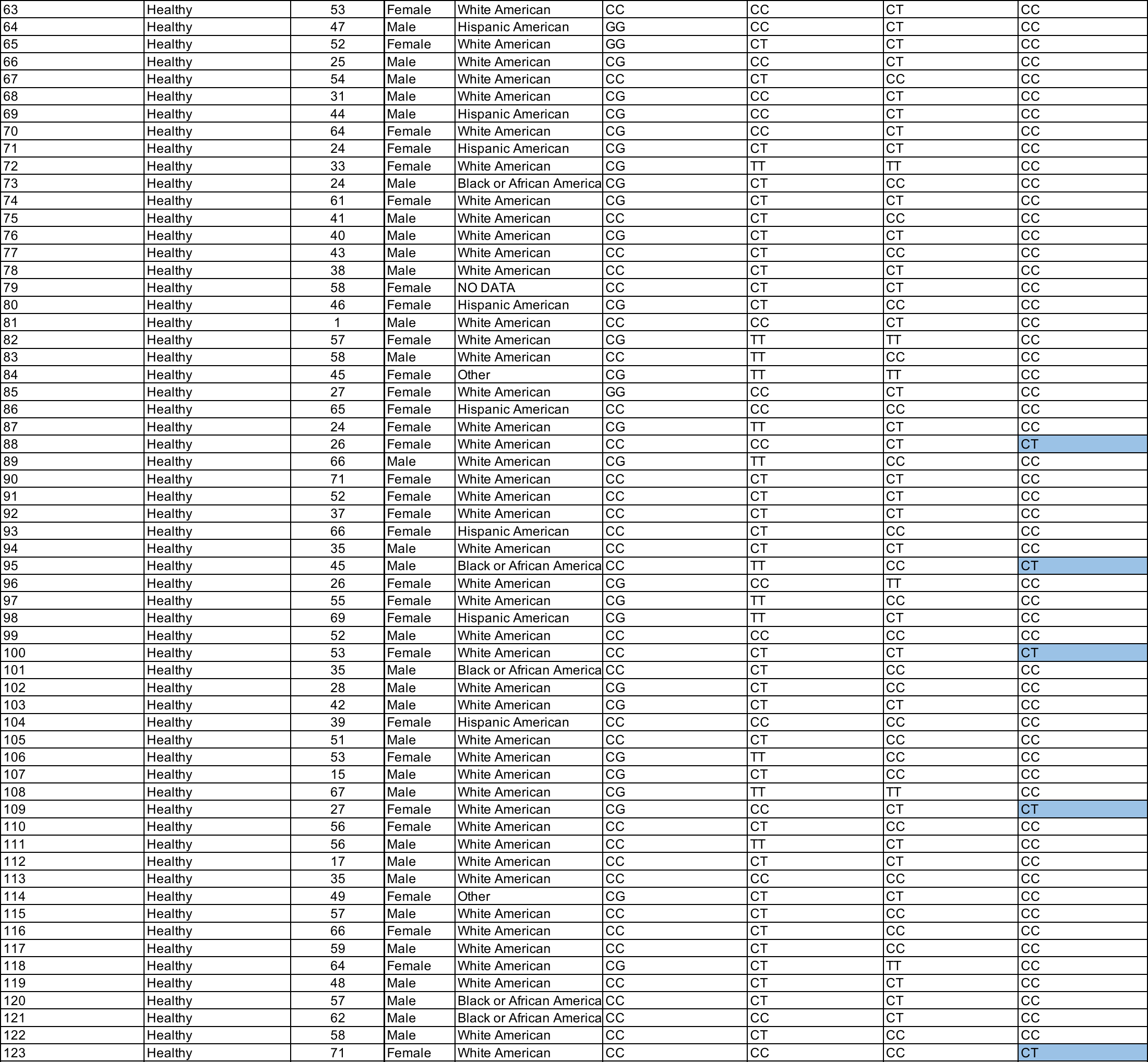

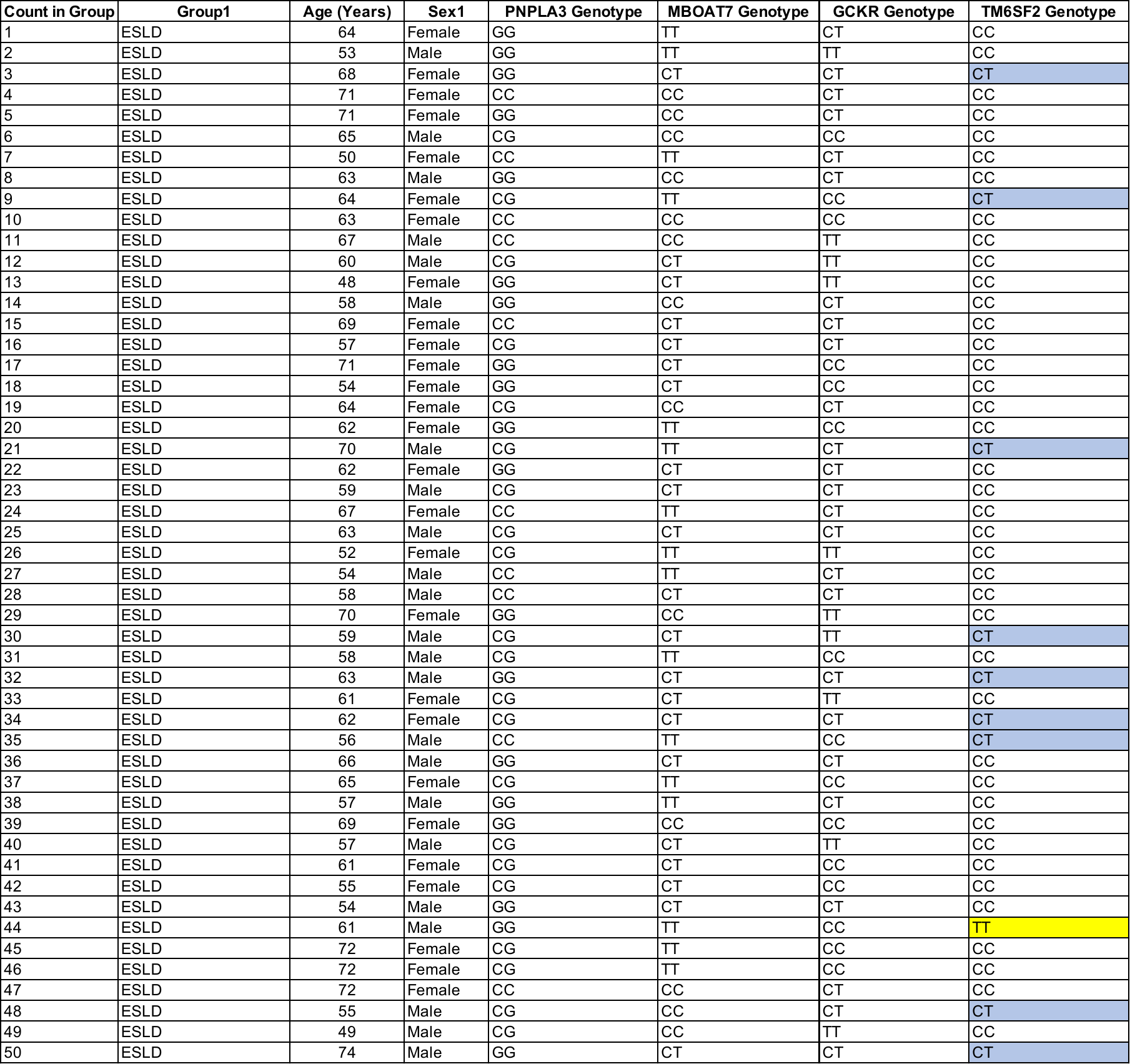
**Supplementary table 5b: Characteristics of ESLD cohorts analyzed.**
